# Autophagy-Dependent Regulation of YAP1 by STK38 Governs Recruitment of Differentiated Cells as Progenitor Cells During Regeneration

**DOI:** 10.1101/2025.04.14.648819

**Authors:** Yongji Zeng, Yang-Zhe Huang, Qing Kay Li, Raymond Ho, Steven J. Bark, Spencer G. Willet, Jason C. Mills

## Abstract

Paligenosis is a conserved cellular plasticity program that allows mature cells to reenter the cell cycle in response to tissue injury. Paligenosis progresses via three stages: autodegradation (with dramatic increase in autophagy and lysosomes), induction of metaplastic or fetal-like genes, and cell cycle entry. Hippo signaling, particularly the downstream effector YAP1, regulates cellular plasticity, but its role in paligenosis has not been studied. Here we first examine paligenosis in digestive-enzyme-secreting chief cells in mouse stomach. We identify Serine/Threonine Kinase 38 (STK38) as a non-canonical YAP1 kinase that phosphorylated and deactivated YAP1 in uninjured chief cells. During paligenosis, STK38 was degraded by autophagy in stage 1, dephosphorylating and activating YAP1. YAP1 activation was necessary and sufficient for the paligenosis that converts chief cells into metaplastic, proliferating progenitors. Additionally, we show STK38, like canonical Hippo kinases, interact with NF2. We also observed the same pattern of YAP1 induction via autophagic destruction of STK38 in other tissues and cell types, suggesting a universal logic model for how the massive autophagy activated in differentiated cells during tissue damage can consequently activate Hippo effectors to induce plasticity for tissue regeneration.

## INTRODUCTION

Paligenosis is an evolutionarily conserved cell plasticity program that mature cells use to reenter the cell cycle.^1^ As such, this program epitomizes the plasticity mature cells can harness in response to tissue injury, allowing organs lacking active stem cells to recruit them when needed.^2^ Paligenosis is characterized by a stepwise process: 1) autodegradation characterized by dramatically increased levels of lysosome and autophagy activation; 2) induction of metaplastic or fetal-like gene expression; and 3) cell cycle reentry.^1^ Given that the need for tissues to recruit stem cells from differentiated cells is nearly universal among multicellular organisms,^3^ and given that conserved cellular changes occur as mature cells reprogram, it is likely that there are also conserved, underlying molecular mechanisms that drive these processes.^4^

The Hippo pathway is broadly conserved and can be a molecular mediator of cell plasticity by inducing wholesale changes in cellular gene expression. YAP1 is the most extensively studied of the downstream Hippo transcriptional regulators. YAP1 is often injury/stress-activated and involved in stem cell activity and induction of a fetal-like repair program in organs as diverse as intestine, liver, skin, and heart.^5–9^ Indirect evidence suggests that YAP1 activation may play a role in paligenosis. For instance, the R-spondin-YAP1 axis promotes *Helicobacter pylori*-induced metaplasia in the stomach^10^ (namely, spasmolytic polypeptide expressing metaplasia, SPEM), and SPEM cells can arise from digestive-enzyme secreting chief cells via paligenosis.^1,2,11^ Targetome analysis has revealed that YAP1 binds to active chromatin elements associated with SPEM-related genes.^12^ Additionally, YAP1 has been implicated in the regulation of autophagy through diverse mechanisms,^13,14^ and upregulation of autophagy is a hallmark of stage 1 of paligenosis.^15^ However, a formal examination of the role for Hippo signaling in paligenosis has not yet been undertaken.

Serine/Threonine Kinase 38 (STK38), also known as Nuclear Dbf2-Related Kinase 1 (NDR1), is a paralog of the canonical Hippo pathway kinases LATS1 and LATS2. Canonical Hippo kinases phosphorylate YAP1, targeting it for proteosomal destruction and blocking downstream YAP1-mediated gene expression changes. STK38 governs a broad range of cellular processes, independent of YAP1; however, despite its homology with LATS1/2, only a few reports have shown that it can also act as a kinase that phosphorylates YAP1.^16,17^ Another key canonical Hippo mediator, NF2 (aka, Merlin), increases LATS1/2 activity and YAP1 destruction by scaffolding the Hippo enzymes to the cytoskeleton. It is not known if NF2 can play an analogous role for a non-canonical Hippo pathway kinase like STK38.

The epithelium of the mammalian stomach body affords a useful model to study both normal, constitutive tissue stem cell function and recruitment of stem cells after injury by paligenosis. Constitutive, non-paligenotic, stem cell activity occurs in a region called the isthmus that is beneath the mucus-secreting pit cells that line the surface of the stomach. Inflammatory injury, like *H. pylori* infection in humans and mice, and toxins like high doses of tamoxifen (HDTAM) in mice, can cause the normally differentiated digestive-enzyme secreting chief cells to undergo paligenosis into proliferative, regenerative SPEM cells.^18–22^ HDTAM causes loss of the secretory-cell maturation transcription factor MIST1 (BHLHA15) within hours, and induces stage 1 of paligenosis along with the accelerated upregulation of autophagy and lysosomes that are dependent on signaling pathways largely mediated by the stress-response transcription factor ATF3.^1,15^

Here, we dissect the intersection between Hippo signaling and paligenosis, mostly using gastric chief cells as an exemplar. We show that YAP1 is both necessary and sufficient for progression through stage 2 and stage 3 of paligenosis. In uninjured chief cells, YAP1 is phosphorylated by STK38 and remains inactive, with chief cells null for *Stk38* and those treated with STK38 antagonists both showing increased YAP1 activation. We show that STK38 can interact with the autophagy machinery via an LC3-interacting region (LIR) and is thus degraded by the upregulation of autophagy in stage 1. As STK38 is degraded, YAP1 becomes activated through dephosphorylation and enters the nucleus to induce targets that promote metaplasia (stage 2 of paligenosis) and proliferation (stage 3). We show STK38 also binds NF2 at homeostasis such that loss of NF2 causes YAP1 activation and induces paligenosis. We show that the degradation of STK38 by autophagy and consequent YAP1 induction occurs in multiple other cell types besides chief cells. In cells that express the secretory-cell-promoting transcription factor MIST1 (BHLHA15), targeting of STK38 autophagic destruction by its LIR domain may be blocked because the LIR domain also binds proteins tagged with the ubiquitin-like protein UFM1. Mature secretory cells maintain elevated MIST1, which in turn directly induces high-abundance UFM1 expression. In paligenosis of secretory cells, MIST1 decrease is an early event,^19,23,24^ so the steady state preservation of STK38 by binding UFMylated proteins would be lost in favor of LC3B interaction and degradation. Overall, the results suggest a universal logic model for recruiting differentiated cells to revert to stem cells: injury induces autophagy, causing degradation of a YAP1 kinase, thereby causing the plasticity-promoting YAP1 to drive paligenosis.

## RESULTS

### YAP1 is activated via de-phosphorylation in stage 1 of paligenosis

To investigate YAP1 regulation and activation, we used the HDTAM injury model^1,25–28^ to induce gastric corpus chief cells of C57BL/6J wild-type mice to undergo paligenosis and adopt a proliferative spasmolytic polypeptide expressing metaplasia (SPEM) phenotype. We performed unbiased phosphoproteomics at 3 timepoints after HDTAM injection: 4 hours (early/pre-stage 1), 12 hours (mid stage 1), and 48 hours (transition from stage 2 to 3). Fig. 1A shows the pattern of YAP1 phosphopeptide abundance during paligenosis. Ser112 in *Mus musculus* (equivalent to human Ser127), is thought to be the principal phosphorylation site that determines YAP1 stability, location, and function.^25,26^ Dephosphorylation at Ser112 activates YAP1 by increasing its stability and allowing it to localize to the nucleus and regulate transcription of target genes.^29,30^ YAP1 peptides phosphorylated on multiple residues from serine 112 to serine 351, including serine 112, all showed markedly decreased abundance at mid-stage 1 (12 hours), with slight rebound for some phosphopeptides by 48 hours (Fig. 1A). Western blotting of a more granular time course showed a similar pattern for phospho-YAP1-S112 with a rebound evident by late stage 3 (72 hours) (Fig 1B). Just as in the phosphoproteomic analysis, phosphorylation in other regions of YAP1 (e.g., Ser382) had differing patterns. For example, a separate phosphorylation site that also leads to YAP1 degradation (pSer382) was not detected at homeostasis or stage 1-2 paligenosis but increased at later timepoints (Fig 1B). Conversely, as expected, antibodies designed to detect activated YAP1 showed approximately the inverse pattern (Fig 1B) although active YAP1 persisted above abundance in uninjured control mice at 72 hours (late stage 3), despite the increase in phosphorylated, inactive forms. We next performed immunofluorescence (IF) to focus on patterns of YAP expression among different gastric corpus cells as well as the intracellular localization of YAP1. Fig. 1C shows YAP1 was activated by 6 hours in chief cells maintaining high abundance through 18 hours and tapering off during transition to stages 2 (starting at 24 hours; note onset of expression of SPEM epitope GS-II) and 3. Active YAP1 was largely confined to nuclei, as expected for it to carry out its transcriptional regulation (Fig. 1C). Immunohistochemistry (IHC) showed that at homeostasis, active YAP1 was largely undetectable in chief cells; however, cells higher in the gastric unit, as has been reported, did express some active YAP1 and showed nuclear localization of YAP1 using an antibody against total YAP1^10,12^ (Extended Data Fig. 1A-B). Thus, the principal changes in YAP1 activation seen in western blotting largely reflected the changes in paligenotic chief cells (because cells higher in the gastric unit express active YAP1 before and after injury; Extended Data Fig. 1A).

**Figure 1.**
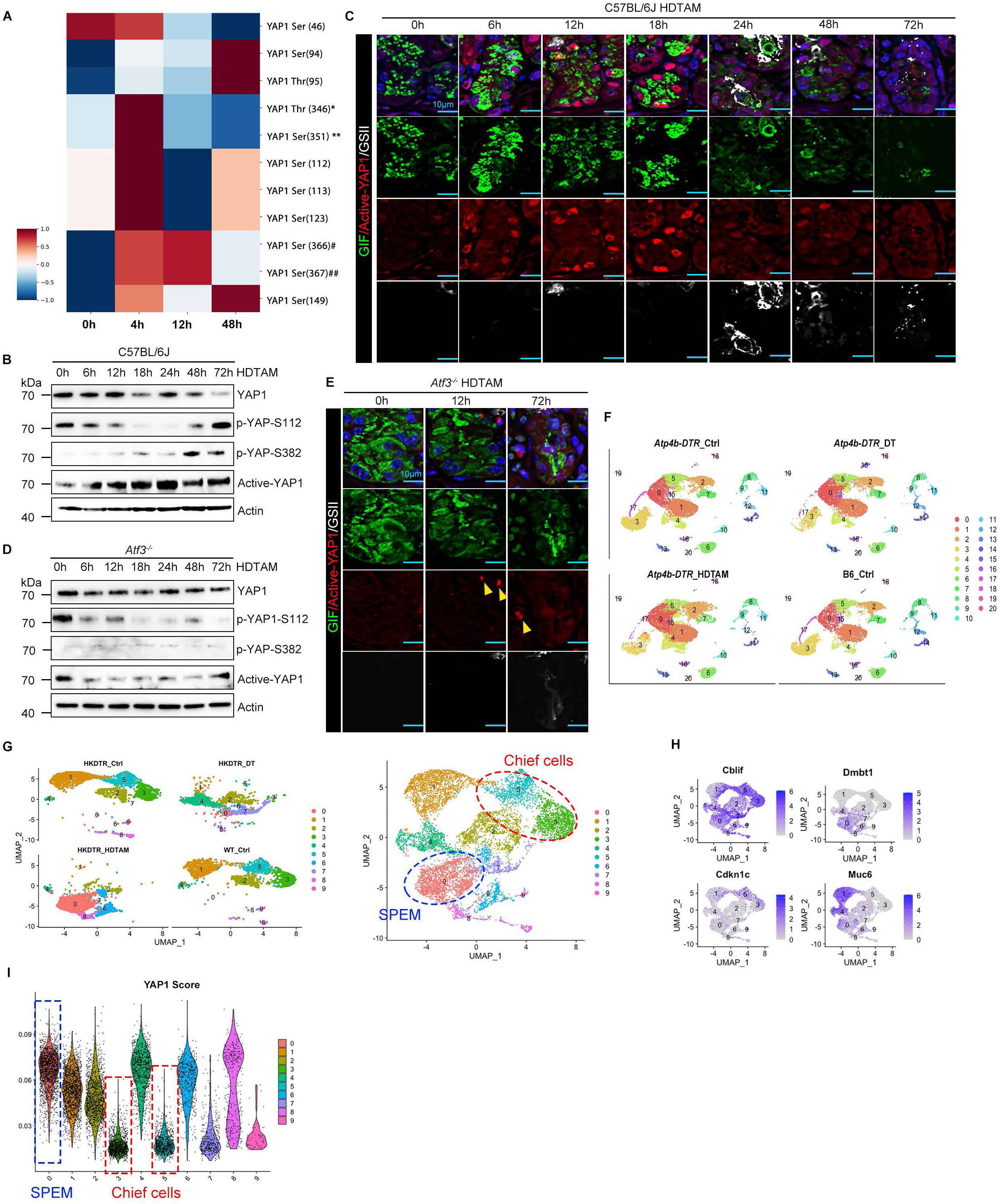
YAP1 is activated via de-phosphorylation in stage 1 of paligenosis. (A) Mass spectrometry-based phosphoproteome profiling reveals dynamic changes in individual YAP1 phosphorylation sites during paligenosis. Phosphorylation sites where the exact residue phosphorylated is ambiguous are marked. Specifically, phosphorylation at * Thr346 could correspond to Thr106, 122, 330, 332, 334, 346, 336, 348, 350, or 352; ** Ser351 could correspond to Ser111, 127, 335, 337, 339, 341, 351, 353, 355, or 357; and # Ser366 could correspond to Ser126, 142, 350, 352, 354, 356, 366, 368, 370, or 372, while ## Ser367 could also correspond to Ser127, 143, 351, 353, 355, 357, 367, 369, 371, or 373. (B) Western blot demonstrates dephosphorylation of YAP1 at site Ser112 during stages 1 and 2 of paligenosis in the HDTAM-treated gastric corpus of C57BL/6J mice. (C) Immunofluorescence reveals less dephosphorylation of YAP1 at site Ser112 (Ser112 in *Mus musculus*, Ser127 in *Homo sapiens*) in chief cells during the early stages of paligenosis. “Active-YAP1” determined by antibody vs dephosphorylated YAP1 Scale bar, 10 µm. (D) Western blot demonstrates no dephosphorylation of YAP1 at site Ser127 during stages 1 and 2 of paligenosis in the HDTAM-treated gastric corpus of *Atf3*^−/−^ mice. (E) Immunofluorescence reveals no increase in YAP lacking phosphorylation at site Ser112 in *Atf3*^−/−^ mice HDTAM-treated chief cells during paligenosis. Yellow arrows point out YAP1-positive mesenchymal cells that are seen at homeostasis and during paligenosis and can serve as immunofluorescence positive controls. (F) Visualization of Uniform Manifold Approximation and Projection (UMAP) plot of single-cell RNA sequencing (scRNA-seq) data from combined gastric corpus cells 48 hours after injury induction with HDTAM and uninjured controls. Twenty-one clusters are annotated in the UMAP plot. (G) UMAP subset clustering of all cells with mucous neck and chief cell lineage markers from Fig.1F (SPEM cells show characteristics of both mucous neck and chief cells.) (H) The mature, non-dividing chief cell marker *Cdkn1c*, aka p57, shows clusters 3 and 5 are mature. Mucous neck cell marker *Muc6* highlights mucous neck cells in clusters 1 and 4. *Cblif* (aka *Gif*) is a chief cell marker also expressed in SPEM. Co-expression of *Muc6* and *Cblif* is characteristic of SPEM and highlights Cluster 0 (and smaller cluster 8) as SPEM-specific, confirmed by SPEM marker *Dmbt1*. (I) Expression of a previously identified gene set of YAP1 downstream target genes shows pronounced expression in SPEM cluster 0 and little expression in mature chief cells. Mucous neck cells show YAP1 expression at baseline, which is consistent with clusters 1 and 4 also showing positivity.

We also took an independent approach to corroborate the findings. We analyzed YAP1 activation by western blot and IF in *Atf3^−/−^* mice, which have profound failure of paligenosis progression with nearly all chief cells dying in transition from stage 1 to 2 (∼18-36 hours^15^). In these mice, which fail to undergo the upregulation of autophagy and lysosomal activity that normally occurs during stage 1, we saw the early loss of pS112 on western blots, but active YAP1 did not emerge. This is consistent with YAP1 activation occurring in stage 1 and during transition into stage 2 of paligenosis (Fig. 1D). As expected, YAP1 was not activated in mutant chief cells (Fig. 1E). As previously reported, some chief cells do survive in *Atf3^−/−^* mice, but they also do not undergo paligenosis. Such remnant chief cells also did not express active YAP1 (Fig. 1E).

To examine a role for YAP1 in transcriptional changes during paligenosis, we performed single-cell RNA sequencing (scRNA-seq) on gastric corpus. This strategy allowed us to focus specifically on paligenotic populations. We examined HDTAM-treated mice with three different control populations. The key control was mice bearing a Atp4b-DTR transgene in which simian diphtheria toxin receptor (DTR) is expressed in gastric parietal cells using elements from a proton pump promoter gene *Atp4b*. Treatment with diphtheria toxin (DT) causes parietal cells to die and activation of normal isthmal stem cells and mucous cells in the neck of the gastric unit, without paligenotic proliferation of chief cells.^31–33^ The other two controls were Atp4b-DTR mice without DT and wild-type B6 mice without HDTAM.

First, we conducted a UMAP analysis on epithelial cells from the four mouse cohorts (Fig. 1F). We then focused on UMAP clustering of mucous neck and chief cell lineages (Fig. 1G). Mucous neck cells are activated to proliferate by loss of parietal cells in both DTR and HDTAM settings, but only chief cells undergo paligenosis to transition into SPEM cells. SPEM cells exhibit a hybrid expression profile of mixed neck and chief cell markers while also expressing unique transcripts.^34,35^ Thus, a UMAP focusing on neck and chief cells would capture our paligenosis-specific population along with other populations activated by injury. Our analysis identified clusters 3 and 5 as mature chief cells, based on well-characterized markers (*Gif/Cblif*, and *Cdkn1c/p57*) (Fig. H). Cluster 1 was likely mucous neck cells based on characteristic markers (e.g., *Muc6*). However, cluster 0 displayed overlapping neck and chief markers (*Cblif* and *Muc6*), consistent with SPEM cells. Moreover, cluster 0 was present almost exclusively in the HDTAM condition and none of the other 3 controls (Figure1G-H). Additional HDTAM-specific clusters (6 and 8) similarly appeared to be secondary SPEM-like populations (Fig. 1G-H).

Next. we generated a “YAP1 score”. We took a set of genes previously shown to be induced by YAP1 *in vivo*.^26^ and used these genes to see which cell populations had highest expression. Clusters 0 and 8 exhibited high YAP1 scores, whereas mature chief cell clusters (3 and 5) had a minimal YAP1 score (Fig. 1I). Together, the findings show that YAP1 is activated in chief cell paligenosis, starting at stage 1 and continuing thereafter to induce a cohort of YAP1 genes in stage 3 (i.e., at the 48h timepoint examined by our scRNA-seq).

Paligenosis is a conserved cellular program across tissues and species, reported in models from *Drosophila* to humans, and from stomach to kidney.^4^ To determine if similar YAP1 patterns could be seen in another well-characterized paligenosis model, we studied the effects of cerulein, an analogue of the hormone CCK, on pancreatic acinar cell paligenosis during acinar-ductal metaplasia (ADM).^1^ We saw a similar pattern of YAP1 activation in late stage 1 (24 hours) with continued YAP1 activity in acinar cells through stage 2 (2-3 days) and into stage 3 (days 5 and beyond) (Extended data Fig. 1C); these results are consistent with other findings showing a role for YAP1 in pancreatic acinar cell metaplasia.^36–38^

### YAP1/TAZ is necessary and sufficient to promote paligenosis in chief cells

To investigate whether YAP1 was required for chief cell paligenosis, we bred mice in which we could conditionally delete YAP1 in chief cells. YAP1 is known as the key effector of kinases in the Hippo pathway, but it also has a paralog, TAZ (aka, WW Domain Containing Transcription Regulator 1, WWTR1), that can perform similar functions in many tissue contexts. Thus, we also targeted TAZ.^39^ Namely, we generated chief cell-specific *Mist1^CreERT^*^2^*^/+^;ROSA26^LSL-Ai^*^9^*^/+^;Yap1^flox/flox^ Taz^flox/flox^* mice. Notably, *Mist1* drives chief-cell-specific expression,^40–42^ and the R26R Rosa locus harbors the Ai9 construct, which is a cre-recombinase-inducible tdTomato allele that lineage traces recombined cells. We used a well-characterized low-dose-tamoxifen-treatment (LDTAM) protocol to induce Cre recombinase in chief cells without causing parietal cell death and chief cell paligenosis.^25–28^ Even the low, baseline nuclear (active) YAP1 in scattered chief cells was reduced specifically in *Yap1^Δ/Δ^*;*Taz^Δ/Δ^* chief cells after LDTAM (Fig. 2A; Extended Data Fig. 2A). However, we did not see any effects on chief cell morphology or census in *Yap1^Δ/Δ^*; *Taz^Δ/Δ^* mature mice even up to 8 weeks after LDTAM (Extended Data Fig. 2B). In contrast, after induction of paligenosis via HDTAM after LDTAM, we noted profound impairment caused by the absence of YAP1 and TAZ. At 48 hours, when most cells transition from stage 2 to 3, we saw significant loss of cells at the base. This phenotype has been called “gland base drop out,” and has been extensively characterized previously as an indicator of failed paligenosis because many cells that fail to undergo paligenosis instead undergo cell death,^43^ particularly in attempted progression to stage 3^1,4,15^ (Fig. 2A,B, Extended Data Fig. 2C,D).

**Figure 2.**
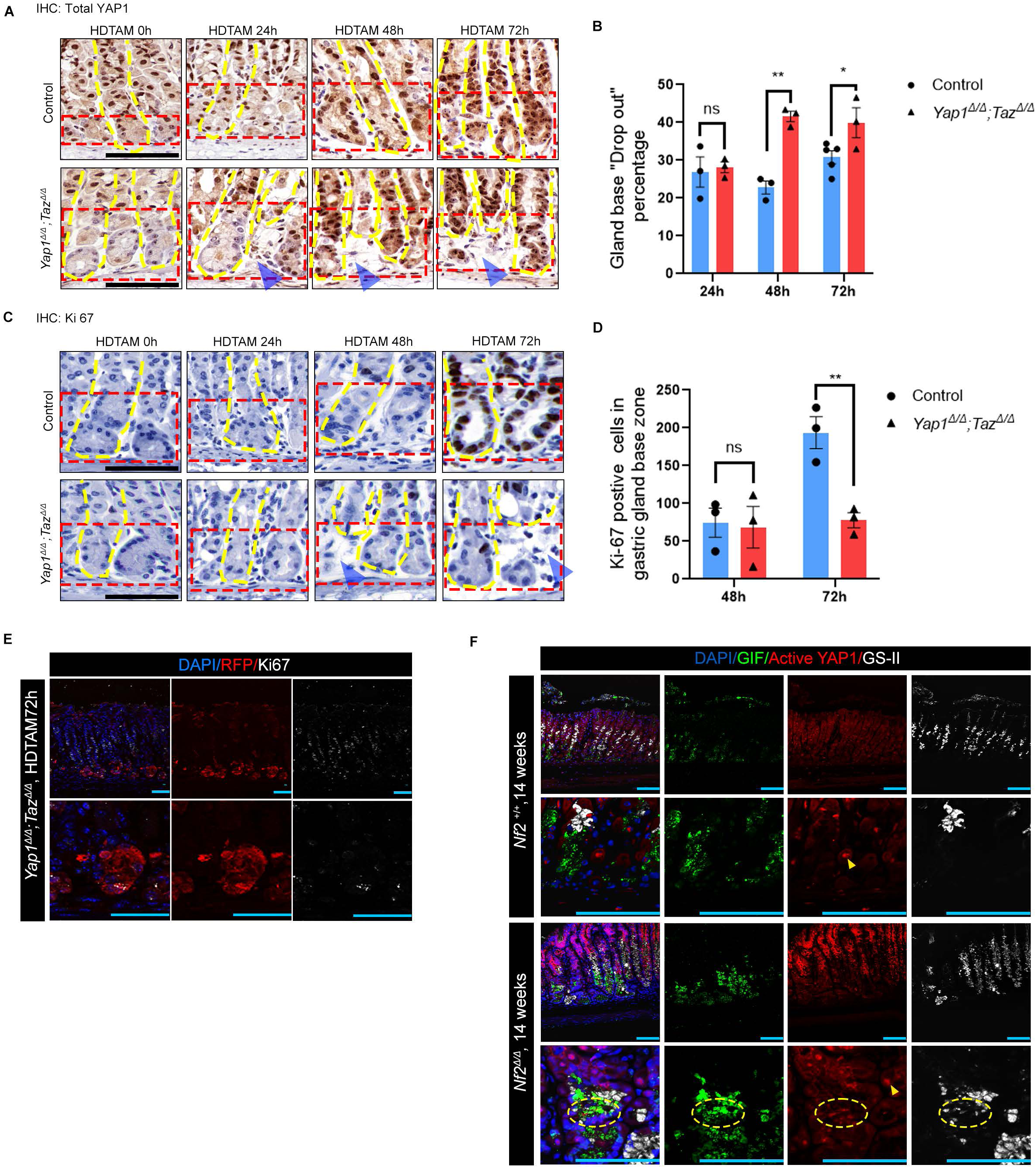
YAP1/TAZ is necessary and sufficient to promote paligenosis of chief cells. (A) Immunohistochemistry (IHC) demonstrates that YAP/TAZ deletion (*Mist1^CreERT^*^2^*^/+^; ROSA26^LSL-Ai^*^9^*^/+^; Yap1^flox/flox^*; *Taz^flox/flox^*) leads to the failure of chief cells to go successfully through the paligenosis process. Top: YAP1 IHC staining of control (*ROSA26^LSL-Ai^*^9^*^/+^; Yap1^flox/flox^*; *Taz^flox/flox^*) stomach during paligenosis. Bottom: YAP1 IHC staining of YAP/TAZ deletion mouse stomach during paligenosis. Red brackets point out the location of the chief cell zone, yellow dashed lines point out the outline of gastric glands, and blue arrows point out the drop out of the gland bases. Scale bar, 50 µm. (B) Quantification of chief cell gland base “Drop out” from Fig. 2A. Note that, due to inefficient recombination of the 4 *Yap1* and *Taz* alleles, many gland bases still have YAP1 and/or TAZ (see YAP1-positive cells at base in panel A), so these quantifications are definitely underestimates. (C) IHC shows YAP/TAZ deletion leads to decreased progression to stage 3 (cell cycle re-entry) in paligenosis. Top: Ki67 IHC staining of control (*ROSA26^LSL-Ai^*^9^*^/+^; Yap1^flox/flox^* ; *Taz^flox/flox^*) stomach during paligenosis. Bottom: Ki67 IHC staining of YAP/TAZ deletion mouse stomach during paligenosis. Red brackets indicate the location of the chief cell zone, yellow dashed lines point out the outline of gastric glands, and blue arrows point out the drop out of the gland bases. Scale bar, 50 µm. “” (E) Immunofluorescence using Ai9 allele to highlight cells in which recombination has occurred due to cre expression from the *Mist1* locus. Note that RFP+, lineage-traced cells are concentrated in the basal chief cell zone, as expected and that these cells, presumably lacking YAP/TAZ, are Ki67-negative (i.e., non proliferating) even well into stage 3 of paligenosis. Scale bar, 50 µm. (F) Immunofluorescence demonstrates the activation of YAP1 in chief cells (*Mist1^CreERT^*^2^*^/+^; ROSA26^LSL-Ai^*^9^*^/+^; Nf2^flox/flox^*) is sufficient to induce SPEM. Yellow arrowheads point out a parietal cell (parietal cells are often YAP1+ at homeostasis) as a positive control for immunolabeling. Yellow dashed region indicates cells at base positive for YAP1, mucous neck cell marker GS-II, and chief cell marker GIF (CBLIF), consistent with SPEM cells. Scale bar, 75 µm. Data information: \*\**P* < 0.01; \*\*\**P* < 0.001 (unpaired t-test). Data are presented as mean ± SEM, derived from 10 low-power fields per condition across three independent experiments.

Cell cycle reentry was also significantly decreased in chief cells, which normally occurs at 48-72 hours (Fig. 2C-D, Extended Data Fig. 2E), confirmed by Ai9 lineage tracing (Fig. 2E). Note that the efficiency of recombination caused by cre recombinase and driven by LDTAM from the *Mist1* promoter is variable, so the quantification of effects on chief cell paligenotic proliferation are likely gross underestimates of the effects of loss of YAP1 and TAZ. Note in Fig. 2A, for instance, how the surviving cells, especially at later stages where cells have already shrunk and entered the cell cycle, all express YAP1, indicating they escaped deletion and were thus able to complete paligenosis. We also analyzed mice with varying combinations of homozygous and heterozygous *Yap1* and *Taz* deletion, and deletion of either gene was sufficient to increase gland-base dropout and significantly decrease paligenotic proliferation (Extended Data Fig. 2F-I).

To investigate whether YAP1/TAZ activation is sufficient to induce chief cell paligenosis, we generated *Mist1^CreERT^*^2^*^/+^; Nf2^flox/flox^; ROSA26^LSL-Ai^*^9^*^/+^* mice. In those mice, the critical Hippo pathway scaffold protein NF2 can be deleted specifically in chief cells. Loss of NF2 is known to decrease Hippo kinase-mediated phosphorylation, thereby leading to activation of YAP1 or TAZ. Accordingly, after LDTAM, we found induced endogenous YAP1 in chief cells (Fig. 2F). The increased YAP1 correlated with a greatly increased fraction of cells in stage 2; i.e., an induction of SPEM cells, as identified by co-expressed GIF (chief cell marker) and GS-II (mucous neck cell marker).^44^ Chronic induction of endogenous YAP1 and TAZ in the *Nf2*^Δ/Δ^ mice disrupted the normal architecture of the gastric unit with expansion of these SPEM cells, many of which were proliferating, indicating increased progression to stage 3 paligenosis (Extended Data Fig. J). Thus, chronic overexpression of endogenous YAP1 activation via *Nf2^Δ/Δ^* was sufficient to induce paligenosis in chief cells. Overall, the findings indicate the activation of the primary Hippo pathway effectors, YAP1/TAZ, is essential and sufficient to drive mature cells into paligenosis.

### STK38 is the only potential upstream kinase of YAP1 whose pattern of activity in paligenosis mirrors the pattern of YAP1 phosphorylation

To determine what regulates YAP1 phosphorylation such that its activity changes in paligenosis, we returned to our phosphoproteomic dataset to identify potential YAP1 upstream kinase(s). Fig. 3A shows all the known YAP1 kinases that were detected in the gastric corpus during the paligenosis time course. Of the canonical Hippo pathway kinases (LATS1 and LATS2, MST1 and MST2), only LATS1 was detected, but neither its abundance nor its phosphorylation state changed in a way consistent with a role in causing the YAP1 phosphorylation pattern during paligenosis (Fig. 3A, Extended Data Fig. 3A). By western blot, we could detect MST1 and (faintly) MST2, neither of which changed in a paligenosis-specific manner. And the total protein abundance of the canonical Hippo pathway YAP1 upstream kinase LATS1, as well as its MST-dependent phosphorylation at Ser908^45^ (Ser909 in *Homo sapiens*), remained unchanged during the paligenosis time course (Fig. 3B). Thus, regulation of YAP1 phosphorylation during paligenosis may be via non-canonical upstream kinases.

**Figure 3.**
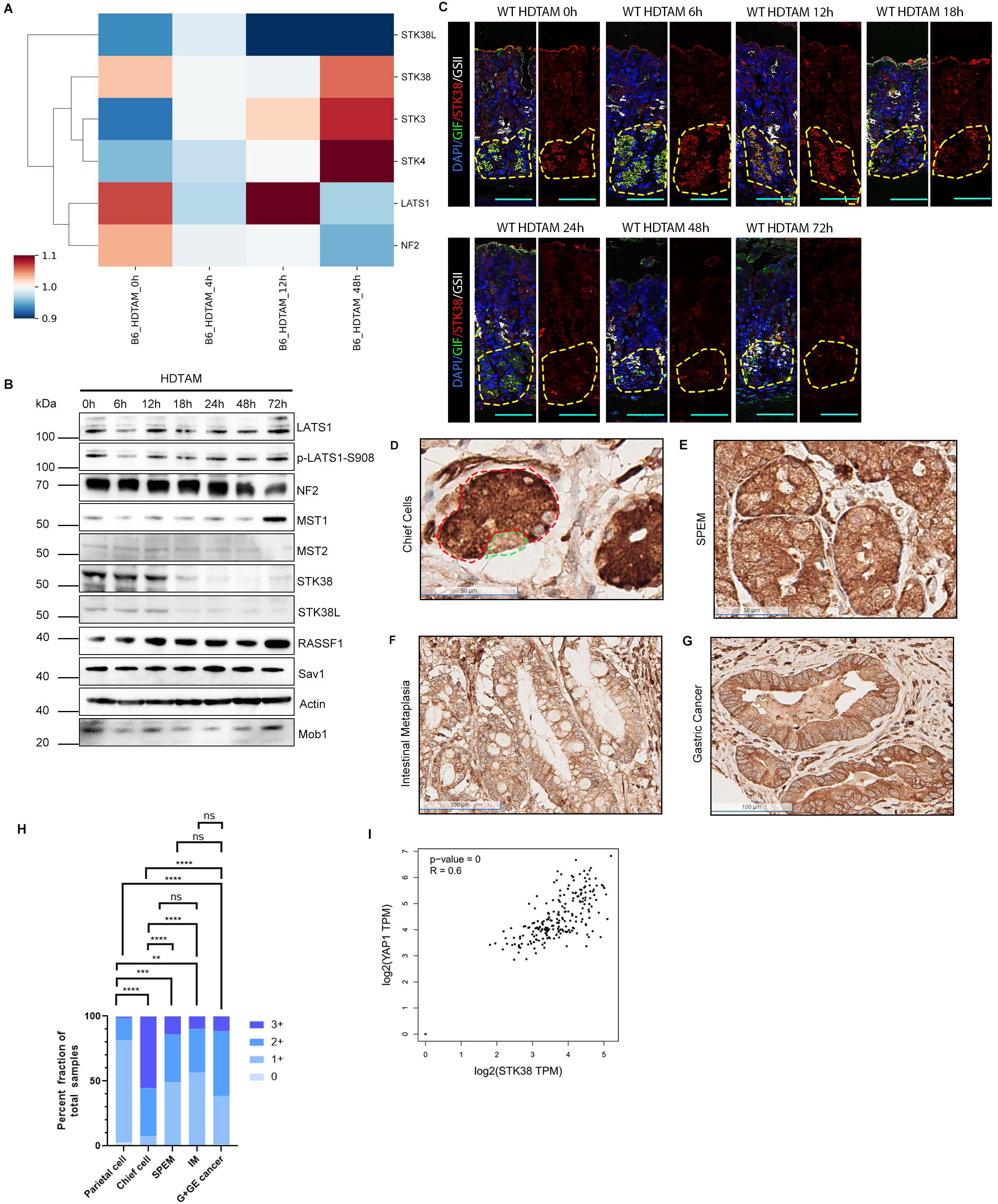
STK38 is the only potential upstream kinase of YAP1 whose pattern of activity in paligenosis mirrors the pattern of YAP1 phosphorylation. (A) Heat map of expression of the principal, known Hippo pathway-related proteins during paligenosis as detected by mass spectrometry. (B) Western blot demonstrates that abundance of the canonical Hippo pathway components (LATS, MST, NF2) remain constant and/or are barely detectable during paligenosis. However, STK38, shows dramatic decrease during stage 1 of paligenosis. (C) Immunofluorescence of wild-type C57BL/6J stomach reveals predominant expression of STK38 in chief cells under homeostatic conditions, with decreasing abundance as paligenosis progresses. Yellow dashed lines outline chief cell zone. Scale bar, 50 µm. D-G: Representative IHC images depicting STK38 protein abundance decrease along the pathway to cancer, from normal human gastric chief cells to Spasmolytic Polypeptide-Expressing Metaplasia, intestinal metaplasia, and gastric adenocarcinoma. (D) Gastric chief cell (red dashed lines) and parietal cell (green dashed lines) in a normal gastric gland. Scale bar, 50 µm. (E) Chief cells undergoing conversion to SPEM (Spasmolytic Polypeptide-Expressing Metaplasia). Scale bar, 50 µm. (F) Intestinal metaplasia. Scale bar, 100 µm. (G) Well-differentiated gastric adenocarcinoma. Scale bar, 100 µm. (H) Significantly down-regulated STK38 protein expression is observed in the progression from SPEM to intestinal metaplasia and gastric or gastroesophageal cancer. \*\**P* < 0.01; \*\*\**P* < 0.001; \*\*\*\**P*< 0.0001 (Fisher’s exact test). *P*-values were calculated by testing for the difference in the mean IHC scores of STK38. (I) Pairwise gene expression correlation analysis using TCGA normal stomach dataset and GTEx stomach dataset reveals a moderate positive linear correlation between *STK38* and *YAP1* expression. (p=8.9e-16, R=0.52, generated by http://gepia.cancer-pku.cn/).

Among the non-canonical Hippo pathway kinases, only STK38 expression correlated with the YAP1 paligenosis phosphorylation pattern by both proteomics and western blot (Fig. 3A,B, Extended Data Fig. 3A). At homeostasis, the STK38 kinase was mainly expressed in chief cells, while during paligenosis, its expression noticeably decreased by 18 hours after HDTAM, correlating with the western blot time course (Fig. 3C, Extended Data Fig. 3B). To determine if the pattern of STK38 abundance across the time course was a conserved aspect of paligenosis, we followed STK38 expression after induction of ADM by injections with cerulein. The reprogramming of mature acinar cells to proliferating ADM cells has been shown to occur via paligenosis.^1,4,15^ As in chief cells, western blotting showed STK38 downregulation began late in stage 1 (24 hours of acinar cell paligenosis) and progressed through stage 2 (48 hours) with little protein remaining by stage 3 (72 hours) (Extended Data Fig. 3C). IHC also showed acinar-specific loss of STK38 by 72 hours (Extended Data Fig. 3D).

Paligenotic progression of chief cells to proliferating SPEM cells also occurs during human pyloric metaplasia. We found that STK38 – just as in mice – was highly expressed in human chief cells and downregulated in SPEM (Fig. 3D). In humans the SPEM that occurs in pyloric metaplasia can progress to intestinal metaplasia (IM) and, eventually, gastric cancer. STK38 remained at low abundance throughout the progression from metaplasia to cancer of both the stomach proper and proximal stomach (gastroesophageal junction) (Fig. 3D-H). In combined TCGA and GTEx gastric expression datasets, we found a moderate positive linear correlation between YAP1 and STK38 at the transcriptional level (Fig. 3J, R=0.6, p<0.001), suggesting that cells that express YAP1 may co-express STK38 to regulate its activity. Overall, both in stomach and pancreas, and in paligenosis and tumorigenesis, STK38, a non-canonical YAP1 kinase, exhibits changes in abundance that are inversely correlated with YAP1 activity.

### STK38 is required for YAP1 activity during paligenosis

To examine whether STK38 is required for YAP1 activation in paligenosis, we next attempted to determine if inhibiting STK38 would induce YAP1. Indeed, mice treated with TAE-684, a STK38 inhibitor, for 7 days showed active YAP1 accumulation in chief cell nuclei (Fig. 4A). We next acquired *Stk38^flox/flox^* sperm, which had been generated as part of the Knockout Mouse Project (KOMP) but not previously characterized, and bred them to generate *Gif^P2A-rtTA/+^;*tetO-Cre*;STK38^flox/fox^* mice, which express cre recombinase to delete functional STK38 in chief cells via rtTA expression from the chief cell-specific *Gif* (*Cblif*) promoter ^46^. Following two weeks of doxycycline induction – and more dramatically after three weeks – active YAP1 began to accumulate in the nuclei of chief cells (Fig. 4B). Western blotting corroborated increased active YAP1 and showed the expected loss of STK38 protein (Fig. 4C). Thus, inhibition of STK38 activity or loss of STK38 is sufficient at homeostasis to activate YAP1 in chief cells.

**Figure 4.**
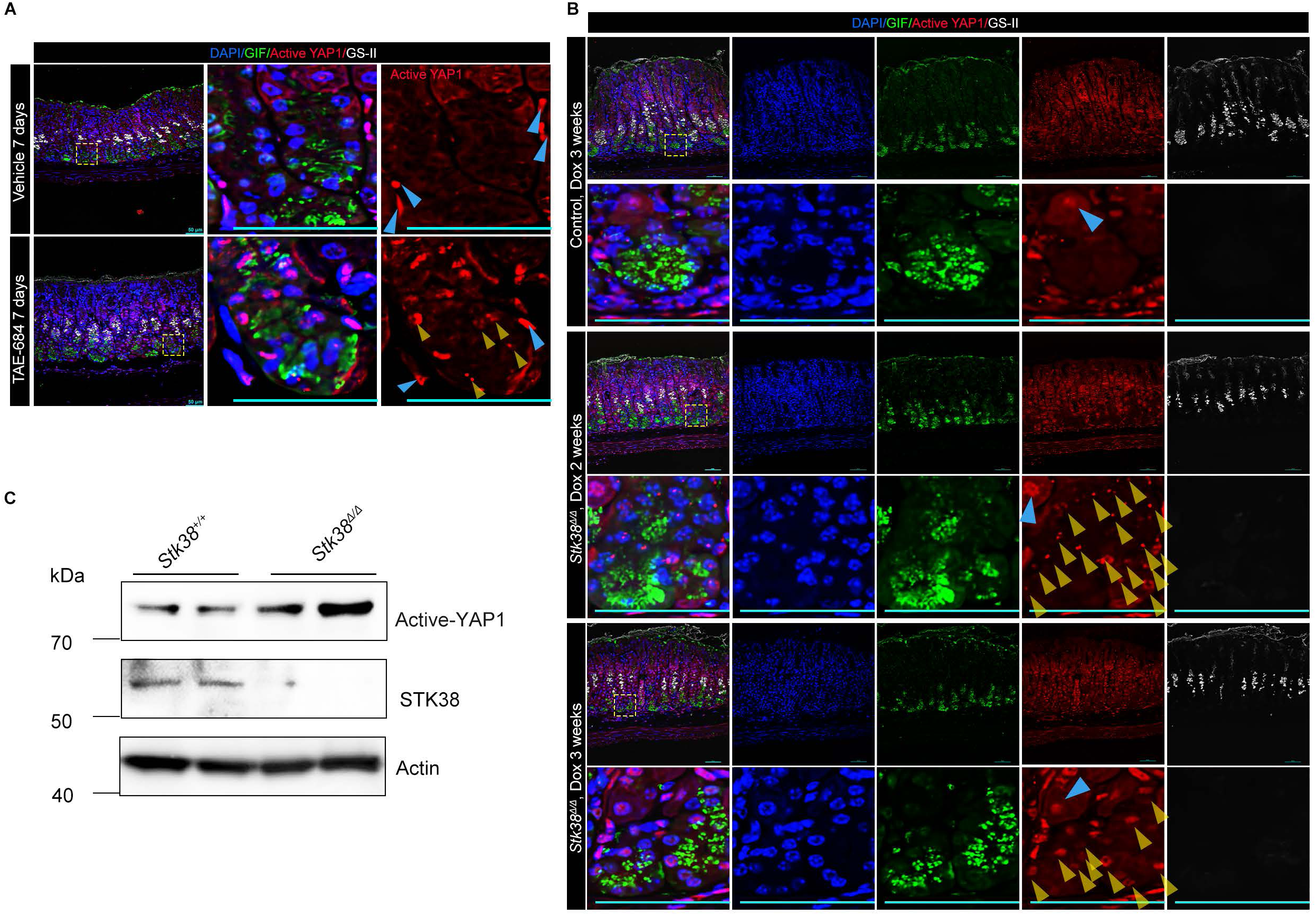
STK38 is required for YAP1 activity during paligenosis. (A) Immunofluorescence of GIF, active YAP1, and GS-II in the stomach of C57BL/6J mice treated with STK38 inhibitor TAE-684. Treatment with inhibitor elevated YAP1 activity in chief cells (yellow arrowheads). Note internal immunostaining control of YAP1-positive mesenchymal cells (blue arrowheads) in both vehicle- and drug-treated tissues. Scale bar, 50 µm. (B) Immunofluorescence demonstrates that loss of *Stk38* in chief cells is sufficient to induce YAP1. Top two rows: control (*tetO-CRE, Stk38^flox/+^*) with 3 weeks Dox treatment; Middle two rows: *Stk38^Δ/Δ^* (*GIF-rtTA, tetO-CRE, Stk38^flox/flox^*), with 2 weeks Dox treatment. Bottom two rows: *Stk38^Δ/Δ^* with 3 weeks Dox treatment. Blue arrowheads indicate parietal cells that can serve as a positive staining control, yellow arrowheads indicate active YAP1 in chief cells. Scale bar, 50 µm. (C) Western blot showing active-YAP1, STK38, and β-actin loading control, indicates that loss of *Stk38* in chief cells is sufficient to induce YAP1 activity in mouse gastric corpus tissue.

### STK38 contains a UFM/LIR site and is lost during transition into stage 2 as it is degraded by autophagy

As shown above, STK38 is the only known YAP1 kinase that is both, expressed in chief cells, and whose expression decreases as YAP1 phosphorylation decreases with YAP1 activity consequently increasing. We next sought to understand the mechanisms driving STK38 decrease in the stage 1 to 2 transition. Previous work has shown that a key feature of paligenosis stage 1 is massive accumulation of autodegradative vesicles (mixes of autophagosomes, lysosomes, autolysosomes).^1,15^ These structures begin to resolve after 24 hours, as cells progress to stage 2. To track these changes on a western blot, we followed abundance of total p62 and p62 phosphorylated at three different sites.^1,47^ p62 began to accumulate at 6 hours, peaked at 18 hours, and started declining at 24 hours (Fig. 5A), consistent with a large accumulation of autophagy-related structures and proteins followed by a period of increased autophagic flux. We also observed a similar accumulation of p62 in pancreatic acinar cells during paligenosis progression to ADM (Extended Data Fig. 4A). During progression to stage 2, flux through the remaining steps of the autophagic program recycles those vesicles, and p62 is degraded (Fig. 5A). ATF3 has been shown to be required for the accumulation of the autodegradative vesicles during paligenosis.^15^ We showed earlier that ATF3 is also required for YAP1 activation (Fig. 1D) and that STK38 abundance decreases during the period of increased autophagic flux (Fig. 3B). Thus, we asked whether the primary mechanism of STK38 loss (and YAP1 activation) was via autophagic flux.

**Figure 5.**
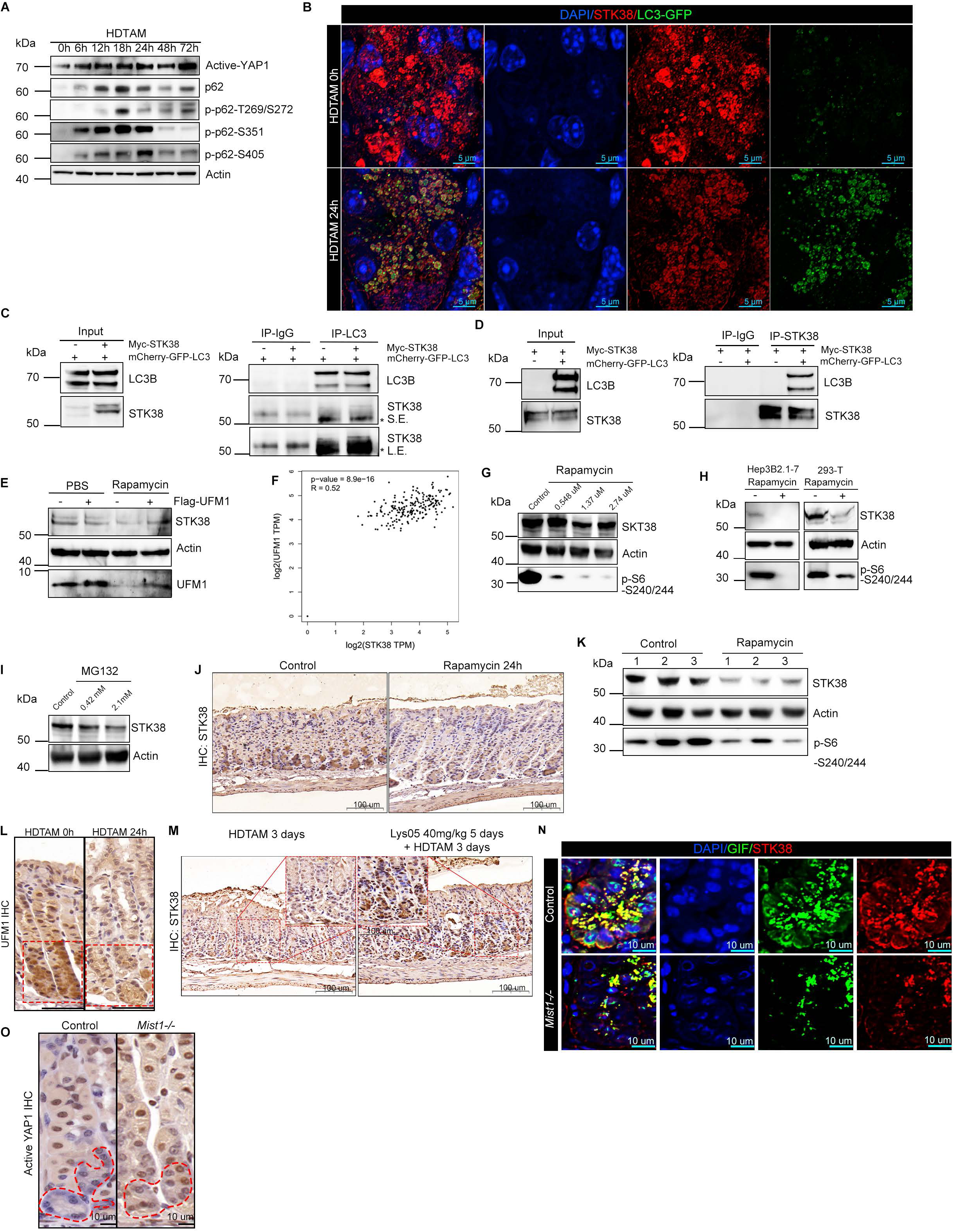
STK38 can bind UFM1 and LC3 and is lost during transition into stage 2 as it is degraded by autophagy. (A) Western blot showing active-YAP1, p62, p-p62T269/272, p-p62-S351, p-p62-S405, and β-actin loading control in whole corpus protein extracts from C57BL/6J mice at various injury time points. (B) Confocal, z-stack immunofluorescence showing STK38 is often found within LC3 vesicles in chief cells in the early stage of paligenosis. Scale bar, 5 µm (C,D) Immunoprecipitation assays demonstrate that overexpressed, myc- and mCherry-GFP-tagged STK38 and LC3 can be co-immunoprecipitated in 293-T cells. “Input” = Total protein lysates of 293-T cells co-transfected with empty vector and mCherry-GFP-LC3 plasmid, or myc-STK38 and mCherry-GFP-LC3 plasmids(C), or empty vector and myc-STK38 plasmid, or myc-STK38 and mCherry-GFP-LC3 plasmids(D). “S.E.” and “L.E.” = short and long exposure. Asterisk denotes molecular mass of tagged STK38, which can be distinguished from immunoglobulin heavy chain seen in input. (E) Western blot demonstrates that overexpression of UMF1 in AGS human gastric cell line is sufficient to protect STK38 from degradation induced by rapamycin treatment. (F) Pairwise gene expression correlation analysis using TCGA normal stomach dataset and GTEx stomach dataset reveals a moderate positive linear correlation between *UFM1* and *STK38* expression. (p=8.9e-16, R=0.52, generated by http://gepia.cancer-pku.cn/). (G,H) Western blot analysis shows that Rapamycin inhibits mTORC1 activity and reduces STK38 protein levels in human (G) AGS (gastric), (H) Hep3B2.1-7 (liver), and 293-T (kidney) cell lines. (I) Western blot shows that MG132 proteasome inhibitor not only does not lead to increased STK38 but results in decrease STK38 protein abundance in a dose-dependent manner in the AGS cell line. (J) IHC of wild-type C57BL/6J stomach tissue demonstrates that rapamycin treatment (60 μg/20 g body weight, euthanized 24hours post-injection) effectively reduces the expression of STK38 in the stomach. (K) Western blot demonstrates that rapamycin treatment effectively reduces STK38 protein expression *in vivo*, as evidenced by gastric protein lysates obtained from experiments like the one depicted in panel J. (L) IHC of wild-type C57BL/6J stomach tissue showing decreased UFM1 in chief cell paligenosis. Red brackets point out the location of the chief cell zone. Scale bar, 50 µm (M) IHC of wild-type C57BL/6J stomach tissue shows that Lys05 treatment effectively preserves STK38 protein levels in HDTAM-treated stomach tissue during stage 3. Scale bar, 100 µm. (N) Immunofluorescence showing reduced STK38 in chief cells (GIF+ cells) of *Mist1*^−/−^ mice compared to control. Scale bar, 10 µm (O) IHC of stomach tissue shows active YAP1 increase in chief cells of *Mist1*^−/−^ mice (right) chief cells compared to control (left). Red dashed lines point out the outline of chief cells. Note YAP1-positive parietal cells serve as positive controls for immunostaining. Scale bar, 10 µm.

Supporting this hypothesis, we observed that in the autodegradation-deficient *Atf3^−/−^* mice, STK38 abundance decreased more slowly and less dramatically compared to wildtype mice (Extended Data Fig. 4B,C), suggesting that STK38 downregulation during paligenosis stage 1 may depend on autodegradation. We next looked in chief cells at homeostasis and at the end of stage 1 to determine if we could see evidence of engulfment of STK38 autophagic machinery. We used GFP-LC3 mice, in which GFP-tagged LC3 highlights membranes of autophagosomes and observed few autophagosomes at homeostasis, as expected, and largescale upregulation of autophagosomes by 24 hours after HDTAM. At 24 hours, STK38 accumulated in LC3-tagged vesicles (i.e., autophagosomes)^48^ (Fig. 5B).

The association of STK38 with LC3 and its localization within autophagosomes indicated STK38 turnover might be regulated by autophagic flux rather than by the more common protein turnover mechanism: proteolysis in the proteosome. We searched for canonical motifs in STK38 that would mediate interaction with LC3 and noticed that both *Mus musculus* and *Homo sapiens* STK38 proteins harbor potential LC3-interacting regions (LIRs). The *Drosophila* ortholog of STK38, Trc, has also been shown to bind ATG8, the ortholog of LC3B^49^. In our *in silico* analysis, using iLIR Autophagy Database (https://ilir.warwick.ac.uk/),^50^ the top scoring potential LIR by Position-Specific Scoring Matrix (PSSM) for the WxxL motif was also a site (CDWWSL) previously shown to bind proteins tagged with UFM1, a ubiquitin-like protein that can be covalently linked to proteins^51,52^ (Extended Data Table 1 and Table 2). Importantly, this proven UFM site and predicted LIR site would allow STK38 interaction with UFMylated proteins to compete for STK38 interaction with LC3B and autophagy machinery, thereby dictating STK38 turnover rate^51^. Specifically, this potential LIR/UFM site would allow STK38 interaction with UFMylated proteins to compete for STK38 interaction with LC3B and autophagy machinery, thereby dictating STK38 turnover rate.

To further explore how LC3B and UFM1 might modulate STK turnover, we turned to a more tractable system to examine molecular interactions: easily-transfectable 293T cells. When we overexpressed both tagged STK38 and tagged LC3 together, we were able to immunoprecipitate STK38 with LC3, and LC3 with STK38, confirming our hypothesis and consistent with orthologous interactions between autophagy machinery and STK38 in *Drosophila* (Fig. 5C,D). Moreover, treatment with rapamycin to induce autophagy in a gastric cell line (AGS cells) decreased endogenous STK38 abundance, consistent with STK38 turnover by autophagy (Fig. 5E). Transfection with UFM1 alleviated the decrease in STK38, consistent with UFM1 binding to STK38 disrupting interactions of STK38 with LC3 and autophagy (Fig. 5E). Analysis of TCGA and GTEx stomach datasets revealed a moderate correlation of *UFM1* and *STK38* transcripts, consistent with both genes being co-expressed (Fig. 5F, R=0.52, p<0.001).

STK38 protein abundance was reduced in a rapamycin dose-dependent manner with phosphorylation of S6 at Ser240/Ser244 used as a positive control to affirm the rapamycin in suppression of mTOR kinase activity (Fig. 5G).^53^ Rapamycin treatment of liver Hep3B2 and kidney 293-T cells showed the same autophagy-dependent degradation of STK38, suggesting that this mechanism for regulating STK38 abundance is not confined to gastric cells (Fig. 5H). Most proteins are degraded by ubiquitination and routing to the proteasome; hence, inhibiting the proteasome increases their abundance. However, AGS cells treated with the proteasome inhibitor, MG-132, actually caused dose-dependent loss of STK38 (Fig. 5I). Loss of STK38 may be because proteasome inhibition leads to induction of autophagy.^54,55^

To test if autophagy regulates STK38 *in vivo*, we injected rapamycin intraperitoneally in mice following a protocol we have used many times to inhibit mTORC1 and induce autophagy.^1,4,56^ The results showed that, even *in vivo*, in mouse stomach and pancreas, autophagy induction caused dramatic loss of STK38 within 24 hours (Fig. 5J,K, Extended Fig. 4D). Conversely, when we blocked autolysosome function with Lys05 during HDT paligenosis, we found that STK38 was preserved even through 72 hours or what is normally a stage 3 timepoint (Fig. 5M). Overall, the data indicate: 1) STK38 downregulation as paligenotic cells progress to stage 2 depends on autophagy, and 2) STK38 binds the autophagy machinery by interacting with LC3 in a manner that could be modulated by STK38 binding UFMylated proteins instead.

### Upstream regulation of STK38 depends on MIST1 induction of UFM1, and on interaction with NF2

We have previously shown that UFM1 is a direct transcriptional target of the transcription factor MIST1 (BHLHA15), which, in the stomach, is specifically expressed in chief cells (as discussed earlier).^57^ Moreover, disruptions of the UFM1 pathway *in vivo* have shown phenotypes in MIST1-expressing cells like plasma cells^57,58^ and Paneth cells.^57,59^ One of the first genes whose expression is decreased in paligenosis is *Mist1*, likely because MIST1 is essential for maintaining the architecture of mature, differentiated secretory cells like gastric chief and pancreatic acinar cells. ^24,60^ Accordingly, UFM1 also decreased during paligenosis (Fig. 5l). As expected, we found that *Mist1^−/−^* mice had decreased UFM1 expression in gastric chief cells and pancreatic acinar cells (Extended data Fig. 4E,F). And, as predicted, STK38 expression was also reduced (Fig. 5N, Extended Data Fig. 4G,H). We were excited to then ask if loss of UFM1 and consequent loss of STK38 would result in constitutive activation of YAP1, as would be predicted by our earlier results. Indeed, *Mist1^−/−^* mice universally showed increased nuclear accumulation of active YAP1 in chief cells. We did not observe a correlation between loss of MIST1 and YAP1 in the pancreas, indicating that additional mechanisms may potentially control STK38 activity and YAP1 activity beyond MIST1-UFM1 in these cells.

Our *in vitro* data indicated that the UFM1-STK38-YAP1 pathway was broadly expressed across organs. As mentioned, a role for both MIST1 and UFM1 has already been reported in plasma cells, so we asked if decreased STK38 and increased YAP1 occurred in plasma cells. Tissue-resident, mature plasma cells are typically strongly MIST1+ and accumulate in tissue like the intervillus compartment in small intestine^58^. Just as in gastric chief cells, *Mist1^−/−^* plasma cells had substantially and significantly increased activated nuclear YAP1 (Fig. 5O, Extended Data Fig. 4I,J). Thus, our evidence indicated a MIST1→UFM1→STK38--|YAP1 axis, with STK38 protein abundance dependent on UFM1 inhibition of its degradation by autophagy, and YAP1 phosphorylation dependent on the outcome of STK38 autophagic degradation. In paligenosis, loss of MIST1 causes loss of UFM1,^57^ which in turn would lead to increased autophagic destruction of STK38 and activation of YAP1.

Our earlier results showed that loss of NF2 led to increased YAP1 activation, which had been as expected based on known stabilization of the canonical Hippo pathway (i.e., LATS and MST kinases) with NF2. However, we also have shown that the canonical Hippo pathway is not the primary regulator of YAP1 in the secretory cells we have been studying. That raised the question of whether STK38 is also stabilized by NF2. Supporting that hypothesis, gastric TCGA and GTEx expression data showed a strong positive linear correlation between *Nf2* and *Stk38* transcripts (Fig. 6A, R=0.72, p<0.001). Moreover, we were able to immunoprecipitate NF2 with STK38 in AGS cells (Fig. 6B). Given that chief cells are only a small fraction of the NF2-expressing cells in the stomach, we decided to examine NF2-STK38 interactions *in vivo* in the pancreas where the large majority of cells are paligenosis-capable, MIST1-expressing acinar cells. Again, as in the *in vitro* studies, we were able to immunoprecipitate NF2 with STK38 (Fig. 6C-E).

**Figure 6.**
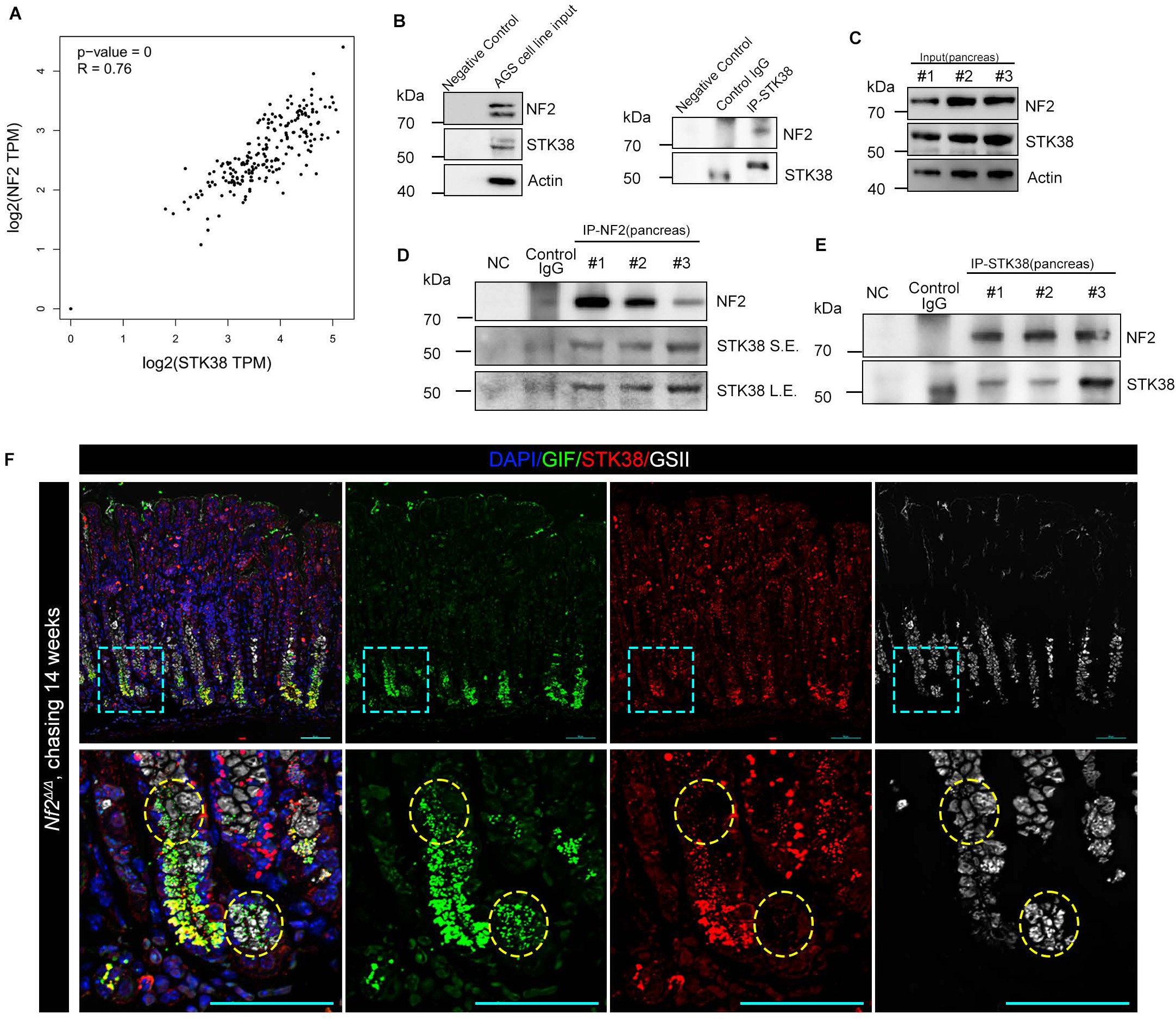
STK38 binds Hippo scaffold NF2. (A) Pairwise gene expression correlation analysis of TCGA and GTEx datasets shows a moderate positive linear correlation between *NF2* and *STK38*. (p=0, R=0.76, generated by http://gepia.cancer-pku.cn/). (B) In AGS human gastric cells, immunoprecipitation of endogenous STK38 co-immunoprecipitates endogenous NF2. C-E: Immunoprecipitation assays demonstrate endogenous NF2 and STK38 protein-protein interactions in the pancreas of C57BL/6J mice. C) Abundance of NF2 and STK38 in protein extract from whole pancreas used as input for immunoprecipitation D) Immunoprecipitation of NF2 co-immunoprecipitates STK38. “L.E., S.E.” = long and short exposures E) Immunoprecipitation of STK38 co-immunoprecipitates NF2 (F) Immunofluorescence for STK38 reveals STK38 downregulation in 8-week-chased *Nf2^Δ/Δ^* mice (*Mist1^CreERT^*^2^*^/+^; ROSA26^LSL-Ai^*^9^*^/+^; Nf^flox/flox^*). Yellow dashed lines highlight the areas of SPEM cells (i.e., GIF/GS-II double-positive cells) resulting from the loss of STK38. Scale bar, 50 µm.

We then returned to the *Mist1^CreERT^*^2^*^/+^; ROSA26^LSL-Ai^*^9^*^/+^; Nf2^flox/flox^* mice, which had shown that loss of NF2 caused YAP1 activation. In those cells that had undergone paligenosis and metaplasia due to loss of NF2 (these are the types of cells in which we found activated YAP1), we observed loss of STK38 as well in cells that showed paligenotic conversion to SPEM cells (Fig. 6F). The results indicate that NF2 interacts with STK38, and its loss leads to reduced STK38 and reduced YAP1 phosphorylation. Thus, NF2 likely regulates YAP1 via its canonical Hippo pathway role in helping to scaffold YAP1 with an upstream kinase, even though the kinase – in this case, STK38 – is not one that is canonically associated with YAP1 (i.e., LATS1/2, MST1/2, the canonical Hippo kinases, are the ones known to be scaffolded by NF2).

## DISCUSSION

Here, we report a signaling axis that dictates Hippo pathway activity in an autophagy-dependent manner in the context of the cellular regeneration program paligenosis. Specifically, we identify STK38 as a non-canonical YAP1 kinase whose turnover is mediated by autophagic degradation. We also show that STK38 targeting to autophagosomes may occur via a STK38 domain that mediates either association with other proteins that are mono-UFMylated or with the autophagy-targeting protein LC3B. Increased UFM1 can help inhibit STK38 autophagic degradation. In many large, protein-secreting cells, UFM1 abundance is increased by the pro-secretory transcription factor MIST1 (BHLHA15). Accordingly, it has previously been shown that *Mist1^−/–^* cells have decreased UFM1 and increased basal autophagy,^57,61^ and we show that they also have more activated (dephosphorylated) YAP1, consistent with decreased STK38 due to continuous autophagic destruction. Additionally, we find that it is not just the canonical Hippo pathway kinases (LATS1 and 2, MST1 and 2) that are bound to NF2, thereby stabilizing their constitutive phosphorylation and destruction of YAP1; as STK38 interacts with NF2 in the same functional way.

As noted, paligenosis is the cellular program mediating return of differentiated cells into the cell cycle via three stages of metabolic and architectural reprogramming. There is evidence that paligenosis is conserved across numerous cell types and species, ^1,2,4,15,61–71^ so it is of significant interest to identify potentially universal regulators of the progression through each stage. The stage 1 to 2 transition depends on a large increase in autophagosomes, lysosomes, and autolysosomes; and genetic or pharmacological inhibition of lysosomal activity blocks paligenosis in stage 1. The stage 2 to 3 transition requires upregulation of mTORC1 activity; and mTORC1 activity is suppressed early in paligenosis by the protein DDIT4 and later by p53.^1,4,65^

Our current results suggest a universal, cell-autonomous logic model for paligenosis (Fig. 7). YAP1 is a highly, evolutionarily conserved protein known to govern cell plasticity in ubiquitous settings.^14,72–74^ A role for YAP1 in paligenosis has not formally been tested, given that the organizing concept of paligenosis has only recently been introduced. However, YAP1 is critical in numerous instances where mature cells are recruited as proliferative progenitors. Some of those YAP1-dependent cellular reprogramming examples, such as recruitment of pancreatic acinar cells, have also been formally shown to occur via the 3-stage paligenosis program.^1,4,15^ Others, such as the reversion to fetal-like state after intestinal injury, have been speculated to occur by paligenosis.^75^ Numerous cases of reprogramming have been shown to depend on initial autophagy/lysosome-requiring steps and later p53/mTORC1-dependent steps (reviewed previously^2,3,76^).

**Figure 7.**
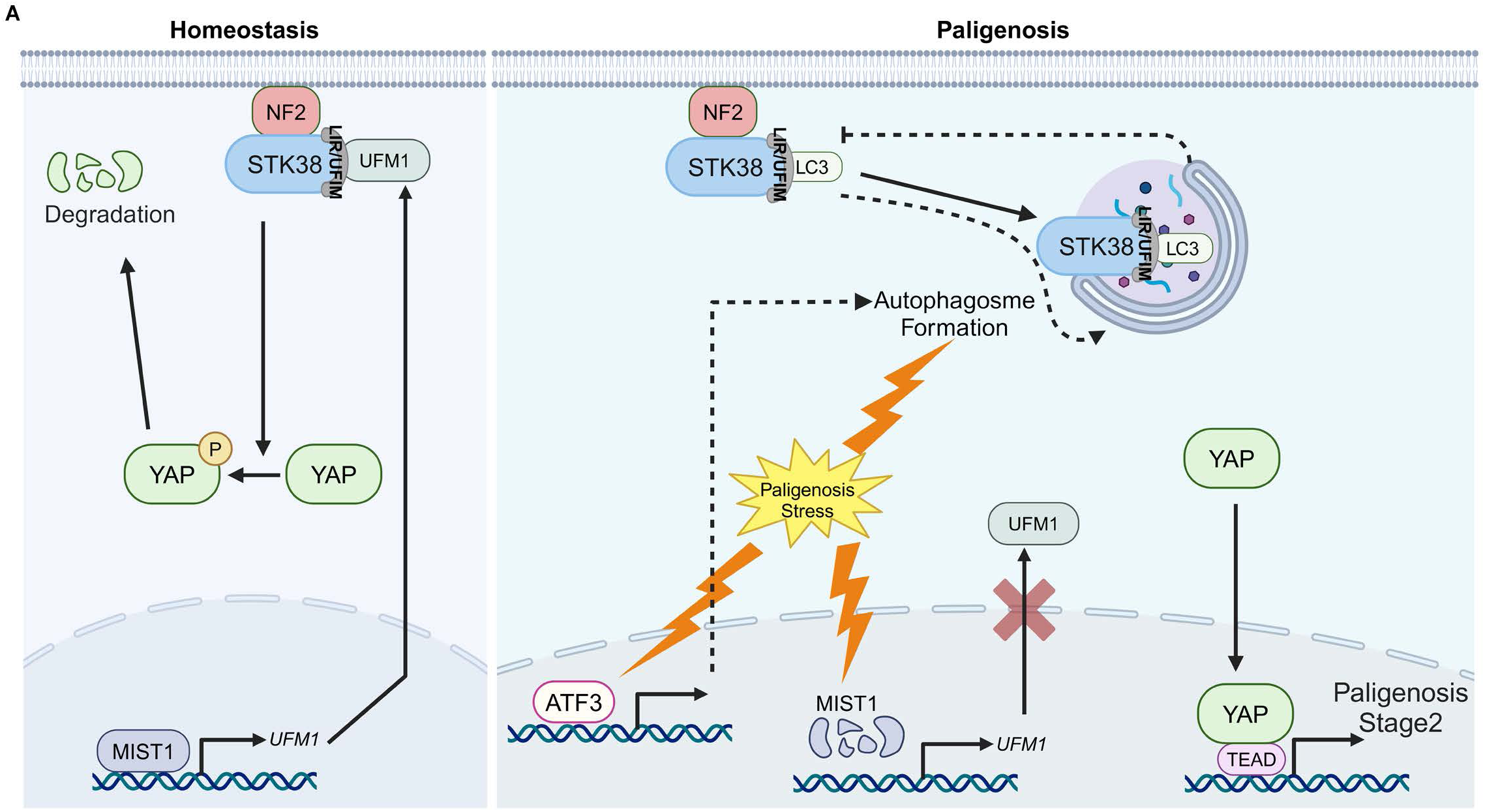
MIST1-UFM1-STK38 axis schematic. (A) Schematic of MIST1-UFM1-STK38 axis regulating YAP1.

Here, we show that YAP1 is required/sufficient for progression to both stage 2 and stage 3. We show how a cell that constitutively expresses both *Yap1* mRNA and an autophagy-sensitive YAP1 kinase can block paligenosis during homeostasis. However, if there is a large-scale cellular insult that leads to activation of massive autodegradation (autophagy/lysosomes), the kinase is destroyed, helping to free YAP1 to induce transcripts that can mediate first a return to a progenitor/fetal-like state (stage 2) and then proliferation (stage 3).

Though our studies hint at a universal paligenotic logic involving YAP1 and autophagy, there may be variation of key components depending on cell type. For example, a potential universal transcription factor that helps induce the autodegradative response is ATF3^15^ as ATF orthologs have similar roles, conserved even to yeast. However, the MIST1-UFM1 axis we describe here may be limited to the large, exocrine secretory cells in which MIST1 dictates secretory architecture.^57,59,60,77–79^ As we show here, plasma cells, which are MIST1-dependent, do have basally increased YAP1 in *Mist1^−/–^* mice. *Mist1^−/–^* Paneth cells (also MIST1-dependent) had a similar phenotype, however, we had difficulty quantifying this difference and so did not report the data here. On the other hand, pancreatic acinar cells, which are MIST1-dependent, did not show basal increased YAP1 in *Mist1^−/–^* mice, although they did show YAP1 dependence on STK38. In short, early loss of MIST1 is almost certainly a generally conserved feature of secretory cell paligenosis (and has already been shown to be such in chief and acinar cells^1,24,80^), because MIST1 promotes mature, post-mitotic secretory architecture and suppresses autophagy; however, which targets of MIST1 are involved in paligenosis (e.g., UFM1) may vary from cell to cell. Moreover, paligenosis occurs in numerous cell types besides MIST1-expressing secretory cells. Thus, the broadest take-home of our studies is that a fundamental feature of paligenosis may be the correlation between injury-induced autophagy that degrades YAP1 kinases, thereby tying the first stage of paligenosis to subsequent stages.

It is unclear how universal STK38’s role is in regulating YAP1 in paligenosis. Few previous studies have implicated STK38 as a principal YAP1 kinase. The Hippo pathway core kinases, LATS1 and LATS2 are proposed to be regulated by two Hippo pathway modules, MST1/2–SAV1–WWC1-3 (HPO1) and MAP4K1-7–NF2 (HPO2).^81^ STK38, like LATS1/2, belongs to the NDR/LATS kinase family and is conserved from yeast to mammals.^17^ STK38 is one of the closest paralogs of LATS kinases and can phosphorylate YAP1 *in vitro.*^17,82^ Moreover, STK38 has been shown to regulate YAP1 phosphorylation in specific tissue types *in vivo*, such as in the intestine, independently of the core Hippo pathway kinase axis (MSTs-LATSs).^16^ In gastric chief cells, LATS1 proteins phosphorylated on Ser909 (Ser908 in *Mus musculus*, activation loop site) and Thr1079 (Thr1078 in *Mus musculus*, hydrophobic motif site) were undetectable at baseline and during paligenosis, and levels of the remaining paralog, STK38L, were also low and unchanged. Thus, we conclude that STK38 is the principal YAP1 kinase in chief cells during paligenosis. Moreover, we surveyed other tissues (pancreatic acinar cells) and cell lines (293T human kidney and Hep3B2 human hepatic) that hinted that the pathway involving autophagy-dependent STK38 turnover, in turn leading to YAP1 dephosphorylation and activation, may be more universal. Considering that ATF3 regulates autophagy partially via Rab7 during paligenosis,^15^ while YAP1 promotes autophagic flux via upregulating RAB7-GAP, ^14^ it makes sense that YAP1 could amplify paligenosis stage1 as progressive loss of STK38 leads to ever more active YAP1 (Fig. 3A-C, Fig. 4A-C). STK38 is broadly expressed and so could regulate YAP1 in many cellular contexts. However, in pancreatic acinar cells MST and LATS may play at least as prominent roles, as they have been shown to mediate Hippo activity in acinar cells, at least during organogenesis, if not in the adult paligenosis context.^83,84^ Interestingly, LATS1 and LATS2 both have potential LIR/UFM motifs that mirror that of STK38. Thus, although we and a previous publication have shown that STK38 can bind UFMylated proteins via an LIR/UFIM domain,^51^ it may be that, in cells where STK38 is not the principal YAP1 paligenosis regulator, LATS may also be protected by homeostatic UFMylated protein binding and degraded by a switch to LC3B interaction at the same motif.

Although we present evidence that MIST1, UFM1, and STK38 all collaborate to modulate YAP1 activation, we expect that other factors also contribute to the sustained YAP1 that helps carry cells through stage 2 and into stage 3. That is because neither loss of MIST1 nor loss of STK38, both of which activated YAP1, were sufficient to induce progression to paligenosis, although forced expression of mutant, degradation-resistant tetO-YAP^S127A^ did do so. One difference might be that YAP1 needs partners, and other injury-induced factors like ATF3 would not be active if YAP1 is induced in the absence of paligenosis-inducing injury. It may be that changes in Wnt are important to pair with YAP1, as an R-spondin-YAP1 axis has been identified that regulates gland regeneration by promoting differentiated cells re-entering the cell cycle when LGR5+ stem cells are lost.^10^ The authors in that case did not test whether this regenerative process occurred by paligenosis, though it seems likely. Another change that occurs when MIST1 is lost (besides decreased UFM1) is restructuring of many aspects of cell architecture, cytoskeletal organization, and cell-cell interaction, ^85^ all of which could ultimately change how cells interact with extracellular matrix (ECM). Given that YAP1 activity is influenced by contact inhibition and mechanotransduction,^86–88^ these additional mechanisms may contribute to YAP1 activation during this stage of paligenosis.

In some ways, the STK38-NF2 interaction is surprising. To our knowledge, this has not been shown previously, but, given the fact that loss of NF2 is sufficient to induce YAP1 activation and eventual paligenosis, and given STK38 is the only known YAP1 kinase operating in chief cells, we had to expect that NF2 works to modulate STK38 activity just as, in its canonical Hippo role, it regulates LATS proteins. But what is somewhat unexpected is that we see most STK38 in the cell cytoplasm, consistent with a baseline interaction with UFMylated proteins, which localize to rough ER,^89–91^ whereas NF2 (Merlin) should be largely on the cell surface. This indicates possibility of intracellular cycling or other yet undescribed mechanisms.

In summary, we have identified a new signaling axis involving STK38, a non-canonical YAP1 kinase, which is regulated by autophagic degradation during the cellular regeneration program paligenosis. This process is likely mediated by the interaction of STK38 with UFMylated proteins protecting against interaction with the autophagy targeting protein LC3B, with increased UFM1 levels inhibiting STK38 degradation (Fig. 5B-5O). Additionally, we demonstrate that STK38 interacts with NF2 in a manner similar to canonical Hippo pathway kinases, further preventing YAP1 activation during homeostasis in secretory cells. Our findings suggest that the MIST1→UFM1 regulation, STK38, and NF2 all collaborate to determine YAP1 activation specifically in secretory cell paligenosis, though the overall theme of YAP1 driving paligenosis, under regulation by an autophagy-sensitive kinase, could be a much more widely conserved logic model for recruitment of mature cells for tissue regeneration.

## Online Methods

### Animal studies and associated reagents

All animal experiments were conducted in accordance with protocols approved by the Institutional Animal Care and Use Committee of Baylor College of Medicine, in compliance with federal regulations. Mice were housed in groups within a specific-pathogen-free barrier facility, maintained on a 12-hour light/dark cycle, and provided with standard chow. Mouse lines were backcrossed to wild-type C57BL/6J (Strain #000664) mice approximately every five generations. The mouse strain used for this research project, LC3-GFP,^92^ *Atp4b-DTR*,^31^ were previously described. B6.129*-Bhlha15^tm^*^3^(cre/ERT2)*^Skz/J^* (Strain #: 029228, referred to as *Mist1^CreERT^*^2^*^/+^*) was a kind gift of Dr. Stephen Konieczny of Purdue University. B6.Cg-*Gt(ROSA)26Sor^tm^*^9^(CAG–tdTomato)*^Hze/J^* (Strain #: 007909, referred to as *ROSA26^LSL-Ai^*^9^) mice were purchased from Jackson Laboratories (Bar Harbor, ME). *Nf2^tm2Gth^* (MGI:1926955, referred to as *Nf2^flox^*) mice were a gift of Prof. Gilles Thomas at INSERM U434, Fondation Jean Dausset, CEPH (Paris, France). Cryopreserved sperm from C57BL/6N*-A^tm1Brd^ Stk38^tm^*^1a^(KOMP)*^Wtsi^/*JMmucd (RRID: MMRRC_049902-UCD, referred to as *STK38^flox^*) mice were obtained from the Mutant Mouse Resource and Research Center (MMRRC) at University of California at Davis, an NIH-funded strain repository, and was donated to the MMRRC by The KOMP Repository, University of California, Davis; originating from Stephen Murray, The Jackson Laboratory. Sperm from these mice were used to fertilize C57BL/6N oocytes for implantation into foster dams by the Baylor College of Medicine Genetically Engineered Rodent Models Core. The *Gif^P2A-rtTA/+^; tetO^Cre^* mouse strain is a kind gift from Dr. Eunyoung Choi, Vanderbilt University School of Medicine (Nashville, TN). The *Yap^flox/flox^* and *Taz^flox/flox^* mice were gifts from the laboratory of Prof. Eric N. Olson at the University of Texas Southwestern Medical Center (Dallas, TX). *Atp4b-DTR* mice were generated at Washington University (St. Louis, MO) as previously described.^29^

Paligenosis was induced by daily intraperitoneal injections of high-dose tamoxifen (HDTAM, 5 mg/20 g body weight; Toronto Research Chemicals) dissolved in 10% ethanol and 90% sunflower oil.^25^ Mice were euthanized at 4 to 72 hours post-injection. Pancreatitis was induced by administering six hourly intraperitoneal injections of cerulein (50 μg/kg in 0.9% saline; SigmaAldrich) every other day for up to one week. To selectively ablate parietal cells, diphtheria toxin (DT, 225 ng/mouse; Sigma) was dissolved in sterile 0.9% sodium chloride saline and administered via intraperitoneal injection once daily for two days. Mice were sacrificed 24 hours after the final injection. Rapamycin (60 μg/20 g body weight; LC Laboratories, R-5000) was administered intraperitoneally in 0.25% Tween-20 and 0.25% polyethylene glycol in phosphate-buffered saline (PBS), and mice were euthanized 24 hours post-injection. Lys05 (40 mg/kg body weight) was intraperitoneally injected two days prior to and during the three daily HDTAM injections. TAE684 (10 mg/kg, Selleckchem, NVP-TAE684) was administered by oral gavage in 30% PEG400, 0.5% Tween 80, 5% propylene glycol, pH 4 in ddH2O once daily for 7 days.

### Immunoprecipitation and western blot

For immunoprecipitation, mouse tissue or cells expressing the specified constructs were collected and lysed in IP lysis buffer (Thermo Fisher Scientific) containing a 1X Protease and Phosphatase Inhibitor Cocktail (Thermo Fisher Scientific). The lysate was then combined with the indicated antibody (refer to Key Resource Table), vortexed briefly, and incubated at 4° C for 1 hour. Pre-washed Protein A/G beads (Santa Cruz Biotechnology) were then added (unless using antibody-conjugated beads), and the mixture was gently rotated at 4° C overnight to allow efficient binding of target proteins to the beads. The next day, bound proteins were separated via the Protein A/G beads, and co-precipitated proteins were eluted for western blot analysis.

For all non-immunoprecipitation western blots, approximately 100 mg of mouse gastric corpus or pancreas tissue was lysed in RIPA lysis and extraction buffer (Thermo Fisher Scientific) supplemented with a 1X protease/phosphatase inhibitor cocktail (Thermo Fisher Scientific). Protein concentration was measured via the BCA protein assay (Thermo Fisher Scientific). Samples were mixed with 5% 2-mercaptoethanol, 1X LDS sample buffer, and then denatured at 95 °C for 10 minutes. Proteins (30 µg) were separated on a 10% NuPAGE 10% Bis-Tris gels and transferred onto nitrocellulose membranes (Invitrogen). Membranes were incubated with 3% bovine serum albumin (BSA) overnight at 4 °C with various primary antibodies: anti-STK38 (1:1,000, Proteintech), anti-STK38L (1:1,000, Proteintech), anti-Phospho-STK38/STK38L (Thr444, Thr442, 1:1,000, Thermo Fisher), anti-active YAP1 (1:1,500, Abcam), anti-Phospho-YAP (Ser127, 1:1,000, CST), anti-Phospho-YAP (Ser397, 1:1,000, CST), anti-YAP (1:2,000, CST), anti-Phospho-SQSTM1/p62 (1:1,000, CST), anti-Phospho-SQSTM1/p62 (Ser403, 1:1,000, CST), anti-Phospho-SQSTM1/p62 (Ser349, 1:1,000, CST), anti-Phospho-SQSTM1/p62 (Thr269/Ser272, 1:1,000, CST), anti-LATS1 (1:1,000, CST), anti-Phospho-LATS1 (Ser909, 1:1,000, CST), anti-Phospho-LATS1 (Thr1079, 1:1,000, CST), anti-NF2 (1:1,000, CST), anti-MST1 (1:1,000, CST), anti-MST2 (1:1,000, CST), anti-Phospho-MST1 (Thr183)/MST2 (Thr180, 1:1,000, CST), anti-MOB1 (1:1,000, CST), anti-SAV1 (1:1,000, CST), anti-RASSF1 (1:1,000, Invitrogen), anti-LC3B (1:2,000, Novus Biologicals), anti-UFM1 (1:1,000, Abcam), anti-Phospho-S6 Ribosomal Protein (Ser240/244, 1:1,000, CST), and anti-beta Actin (1:1,000, Santa Cruz). Membranes were then incubated with HRP-linked secondary antibodies (CST). Protein intensities were normalized to a tubulin loading control, with chemiluminescent signal quantified using ImageJ under non-saturating conditions.

### Immunofluorescence and immunohistochemistry

For immunofluorescence, mouse tissues were excised and flushed with PBS, then inflated with freshly prepared 3.7% formaldehyde in PBS (i.e., freshly made formalin). The stomach was clamped, suspended in fixative within a 50-mL conical tube overnight, then rinsed in PBS followed by 70% ethanol. Samples were arranged in 3% agar within a tissue cassette and processed for paraffin embedding. Sections (5 μm) were cut, deparaffinized, and rehydrated through graded Histo-Clear (National Diagnostics), ethanol, and water. Antigen retrieval was conducted in sodium citrate buffer (2.94 g sodium citrate, 500 μL Tween 20, pH 6.0) using a pressure cooker. Slides were blocked with 5% normal serum and 0.2% Triton X-100 in PBS. Primary antibodies (see Key Resources Table) were applied overnight at 4° C, followed by PBS rinsing, a 1-hour room-temperature incubation with secondary antibodies and/or fluorescently labeled lectin, additional PBS rinsing, and mounting with ProLong Gold antifade mountant with DAPI (Molecular Probes).

For immunohistochemistry (IHC), the protocol followed the immunofluorescence procedure with the following modifications. Slides were first deparaffinized through serial Histoclear washes and rehydrated in an ethanol series. A 15-minute quenching step in methanol containing 1.5% hydrogen peroxide was added before heat-mediated antigen retrieval. After applying the secondary antibody, color development was achieved using the Vectastain Elite ABC HRP Kit (Peroxidase, Standard) (Vector Laboratories) according to the manufacturer’s instructions. Slides were then developed with the DAB Substrate Kit (Thermo Scientific) and mounted in Permount Mounting Medium (Fisher Chemical).

### Imaging

Immunofluorescence microscopy images were acquired using the AX R Confocal System with an Eclipse Ti2-E Inverted Microscope (Nikon), maintaining consistent exposure times across samples for accurate comparison. Brightfield images were acquired on an Epredia 3D Histech Pannoramic MIDI II brightfield slide scanner. Image analysis and post-processing adjustments were completed using Adobe Photoshop 2024.

### Proteomics

Approximately 100 mg of mouse corpus stomach tissue for each mouse was submitted to Baylor College of Medicine Proteomics Core Facility for further proteomics analysis. The proteomics data was obtained from the Baylor College of Medicine Proteomics Core Facility. The experimental protein mass spectrometry data was compared to a liver reference mixture (RefMix) of proteins using a TMT mass tagging approach (iBAQ). These data were reported as an Excel file containing protein identifiers (ENSEMBL IDs and gene symbols), spectral counts, and iBAQ area comparisons for each protein identified in the experiment. For the analysis reported here, the experimental iBAQ area data for each identified protein at each time point was normalized using the RefMix for the same protein in the same experiment. These normalized data were then converted to a Z-score across the sample time points. Clustered heatmaps were generated based on these Z-score comparisons using Python libraries Pandas (version=2.0.3), Matplotlib (version=3.8.0), and Seaborn (version=0.12.2). The processed proteomics and phosphor-proteomics-site data spreadsheets are in Extended Table 3 and Extended Table 4.

### Single cell RNA-seq tissue preparation and processing

Single cell RNA-seq tissue preparation and processing have been previously described.^29^ Fresh gastric corpus tissue was collected from both vehicle (uninjured), DT-treated and HDTAM-treated mice. After flushing with ice-cold PBS and removing the forestomach and antrum, the tissue was washed in cold DMEM. For mechanical dissociation, the corpus was shaken in pre-warmed digestion solution (Advanced DMEM/F12 with HEPES, BSA, antibiotics, gentamicin, and collagenase) at 37° C for 30 minutes. Solid stomach musculature was discarded, and the cell suspension was washed in rinse media before chemical dissociation with 0.05% Trypsin-EDTA at 37° C for 5–15 minutes, with occasional mixing. Cell viability and single-cell dissociation were monitored with Trypan blue staining, and cell counts were taken using a hemocytometer. Upon reaching single-cell suspension, digestion was neutralized with DMEM + 10% FBS, filtered with a 30 µm filter, and submitted to Baylor College of Medicine Single Cell Genomics Core.

Tissue for scRNA-seq was processed by Baylor College of Medicine Single Cell Genomics Core using the 10x Genomics platform. Raw data from control and HDTAM 48-hour samples were processed with CellRanger 6.1.2 using the mouse reference genome mm10. Data were quality controlled in Seurat/R to filter out doublets and low-quality cells based on mitochondrial content (mito.percent < 0.5) and read thresholds (nFeature > 700 and nCount < 75000), and then globally scaled and normalized. PCA was conducted on the top 2,000 variable genes, followed by UMAP for unbiased clustering with dims = 1:23 and resolution = 0.3. Clusters were annotated using known cell markers from previous studies in the laboratory, as well as with reference gastric single-cell atlases. The analysis code is available on FigShare. RNA-seq data have been deposited in Gene Expression Omnibus with accession # ___.

### Cell culture

AGS cells (ATCC) were maintained in Advanced DMEM/F12 with 10% FBS, nonessential amino acids, and 1% Pen/Strep at 37 °C and 5% CO₂. Transfection of AGS cells with the pRK5-FLAG-UFM1 plasmid (Addgene) was performed using Lipofectamine 3000 (Thermo Fisher Scientific) according to the manufacturer’s instructions, followed by treatment with 2.74 μM rapamycin for 24 hours.

293T cells (ATCC) were cultured in DMEM with 10% FBS, nonessential amino acids, and 1% Pen/Strep at 37 °C and 5% CO₂. For overexpression studies, 293T cells were transfected with pFLAG-YAP1 (Addgene), pClneoMyc human STK38 (Addgene), pRK5-FLAG-UFM1 (Addgene) using Attractene Transfection Reagent (Qiagen), following the recommended protocol.

Hep 3B2.1-7 cells (ATCC) were cultured in ATCC-formulated Eagle’s Minimum Essential Medium with 10% FBS, and 1% Pen/Strep at 37 °C and 5% CO₂.

### Human tissue microarray studies

The studies utilized human tissue microarrays (TMA) constructed from formalin-fixed pathology material. The use of formalin-fixed pathology samples was reviewed and approved by the Institutional Review Board of Johns Hopkins Medical Institutions. Written informed consent for participation was not required for this study in accordance with the national legislation and the institutional requirements. Study case collection, construction of the TMAs, immunohistochemical staining of markers, evaluation and scoring of IHC staining, data analyses and statistics have been previously described.^93^

### Expression constructs

The following plasmids were purchased from Addgene: pClneoMyc human NDR1 (#37023), pFLAG-YAP1 (#66853), pRK5-FLAG-UFM1(#34640).

### Quantification and statistical analysis

Ki67+ proliferative cells were quantified by counting at least 90 fully integrated glands from multiple randomly selected regions of the stomach corpus, unless otherwise noted in the manuscript. Active-YAP1+ plasma cells were quantified by counting at least 100 CD138+ plasma cells from multiple randomly selected regions of the intestine villus. Chief cell “drop out” at the gastric gland base during HDTAM injury and recovery was assessed based on morphology in H&E-stained sections, as previously described.^43^ Sample sizes were determined through power analysis to ensure statistical significance while minimizing the number of mice used. For statistical analysis, all time points and treatments included at least three mice of both sexes, unless specified in the results or figure legends. Outliers were not excluded from analysis. Student’s t-test was used for comparisons between two conditions when assumptions were met. For human tissue microarray analysis, differential expression of STK38 across phenotypic groups was assessed by Fisher’s exact test. All analysis and graphing were performed using Prism (GraphPad), with a P-value < 0.05 considered significant. Statistical significance is indicated as (*), (**), (***) or (****) for P-values < 0.05, 0.005, 0.0005, and 0.0001, respectively, and “n.s.” denotes not significant. Samples were randomized, and measurements were blinded to minimize experimental bias.

## Data availability

The original single cell RNA sequence data are available at GEO (GSE294328).

## Author contributions

Y.Z. designed and performed experiments, analyzed data, conducted bioinformatics analyses, provided funding, wrote, and edited the manuscript. Y.-Z.H. and S.J.B conducted bioinformatics analysis. Q.K.L., R.H. and S.G.W conducted experiments and analyzed data. Q.K.L. accrued and analyzed human tissue specimens. J.C.M. analyzed data, conducted bioinformatics analyses, provided funding, and wrote and edited the manuscript. All authors provided feedback for the manuscript.

## Acknowledgements

We thank Dr. Joseph Burclaff for gastric tissue slides from Lys05-treated mice. We also thank Dr. Eunyoung Choi from Vanderbilt University School of Medicine (Nashville, TN) for providing us with the *Gif^P2A-rtTA/+^; tetO^Cre^* mouse strain. We thank Pamela Parsons and the Texas Medical Center Digestive Disease Center Tissue Analysis and Molecular Imaging Core (supported by NIH, P30DK056338) for their assistance with tissue preparation for histology. We thank the Baylor College of Medicine Mass Spectrometry Proteomics Core (supported by NIH, P30CA125123, P50HD103555, P50HD103555, S10OD026804; Cancer Prevention and Research Institute of Texas, RP170005, RP210227) for helping us to produce the proteomics raw data. We thank Baylor College of Medicine Genomic and RNA Profiling Core, also part of the Texas Medical Center Digestive Disease Center FGM Core (supported by NIH, P30DK056338, P30CA125123, P30ES030285, S10OD036427; Cancer Prevention and Research Institute of Texas, RP200504) for helping us to produce single cell RNA sequencing raw data. We thank the Baylor College of Medicine Genetically Engineered Rodent Models Core. We also extend our special thanks to Dr. Robert M. Lawrence for his editorial suggestions during the writing and figure preparations. The graphical abstract was created using BioRender.com. The following authors of this work were supported by the following grants: Y.Z., National Cancer Center Postdoctoral Fellowship; J.C.M., NIH R01DK094989, R01DK105129, R01CA239645, R01CA246208, and R01DK134531.

## Supplemental Figure Legend

**Extended Data Figure 1.**
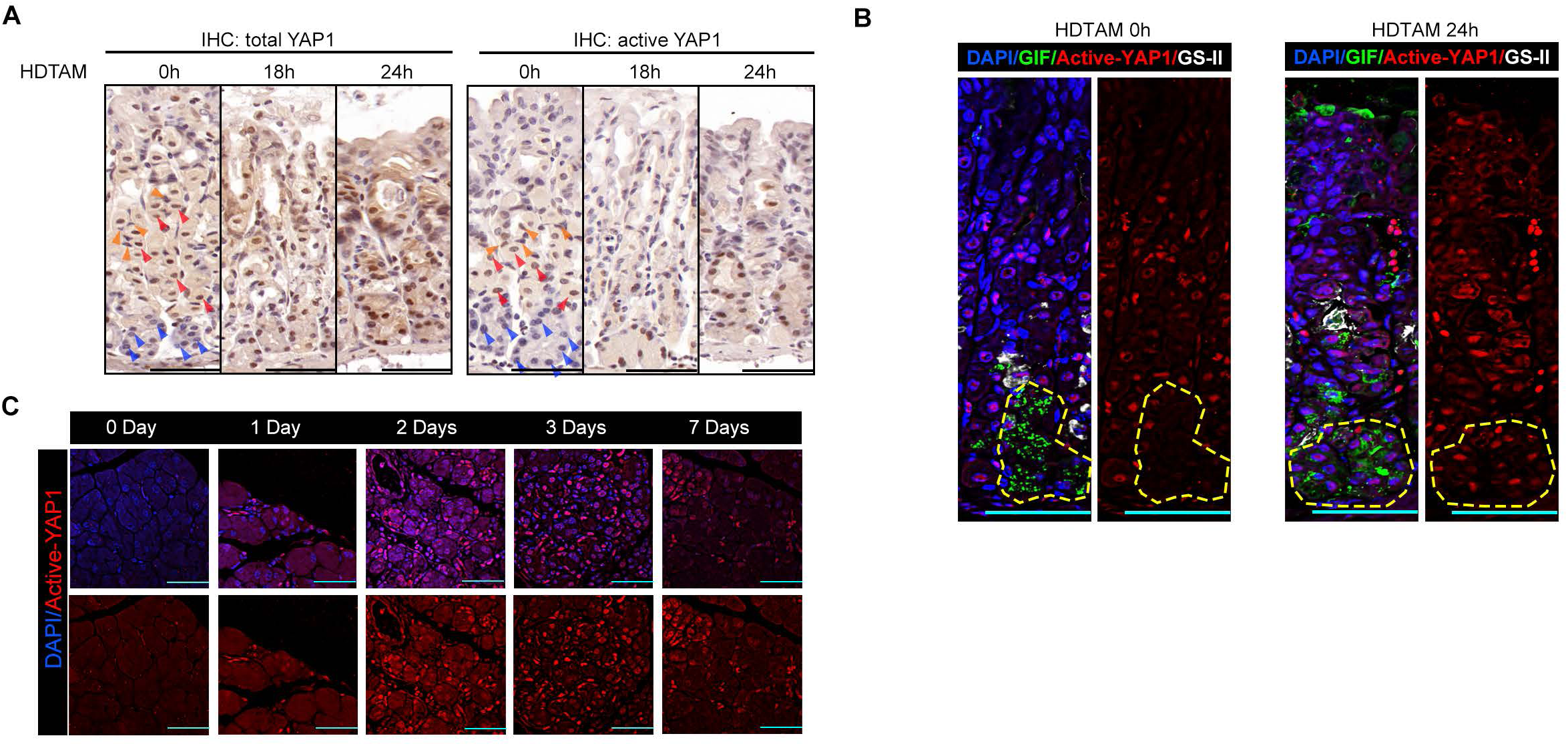
YAP1 activation via dephosphorylation occurs by late stage 1 of paligenosis. (A) Immunofluorescence for total YAP1 (left) and active YAP1 (dephosphorylation at site Ser112) (right) in mouse gastric glands during homeostasis (0 h after HDTAM) and stage 1 paligenosis (18 h and 24 h after HDTAM) in HDTAM-treated C57BL/6J mice. The blue arrowheads, red arrowheads, and orange arrowheads indicate examples of chief, parietal, and mucous neck cells in the gastric gland, respectively. Note that parietal and neck cells express some nuclear total YAP1 and active YAP1 at homeostasis, but chief cells only have active, nuclear YAP1 during paligenosis. (B) Immunofluorescence staining of active YAP1 in chief cells during homeostasis (left) and late stage 1 of paligenosis (right) in HDTAM-treated C57BL/6J mice. The yellow cycle area indicates the chief cell zone of the gastric gland. (C) Elevated levels of active YAP1 observed in the pancreas of cerulein-treated C57BL/6J mice.

**Extended Data Figure 2.**
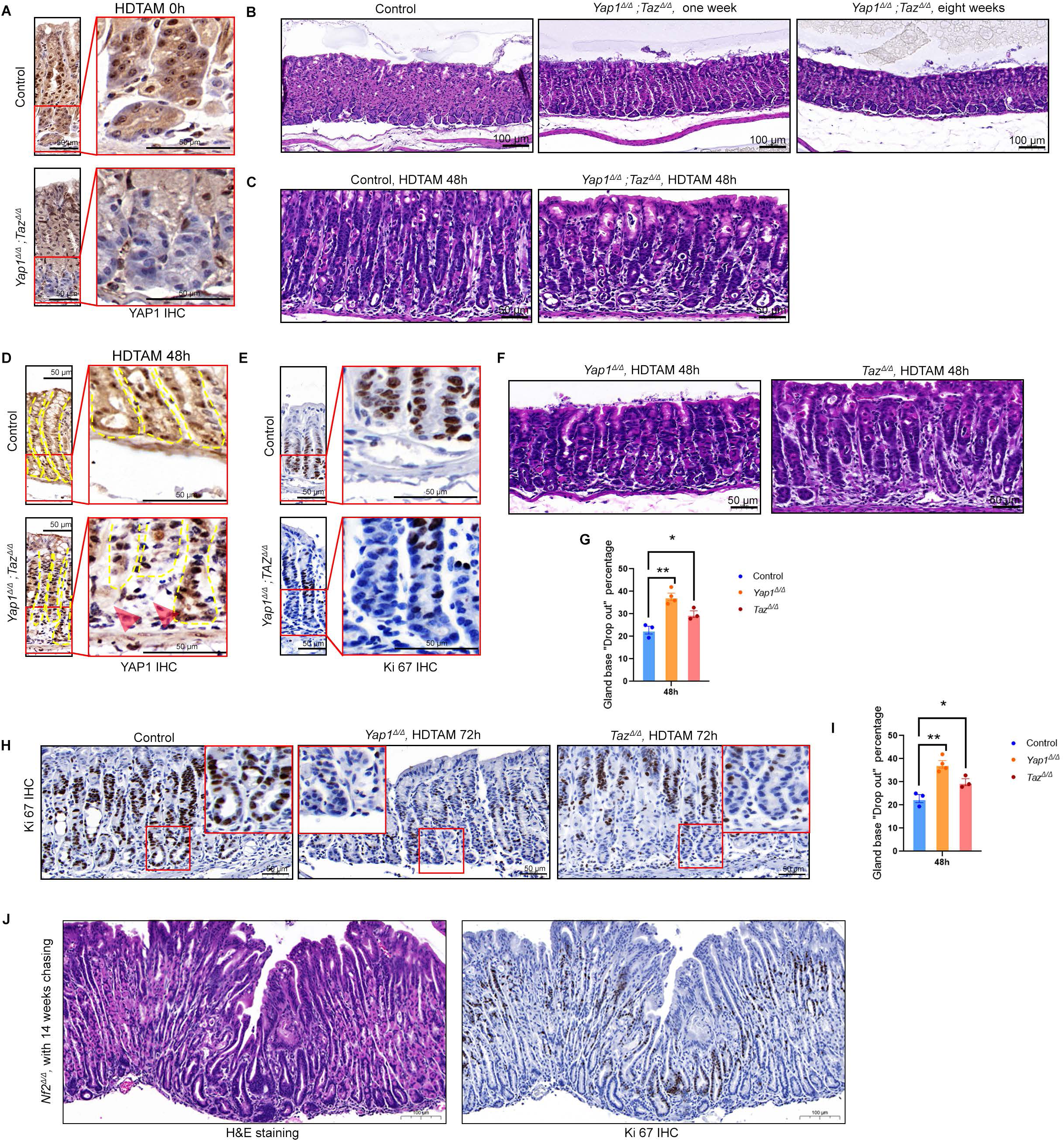
Hippo pathway downstream effectors YAP1 and TAZ are necessary and sufficient to induce paligenosis. (A) YAP1 IHC shows the efficiency of chief cell YAP1 deletion in *Yap^Δ/Δ^; Taz^Δ/Δ^*(*Mist1^CreERT^*^2^*^/+^; ROSA26^LSL-Ai^*^9^*^/+^; Yap1^flox/flox^* ; *Taz^flox/flox^*) mouse gastric gland. Top: YAP1 in control (*ROSA26^LSL-Ai^*^9^*^/+^; Yap1^flox/flox^*; *Taz^flox/flox^*) stomach under homeostasis. Bottom: YAP1 IHC staining of YAP/TAZ deletion mouse stomach under homeostasis. Red brackets point out the location of the chief cell zone. Scale bar, 50 µm. (B) H&E images show that *Yap^Δ/Δ^; Taz^Δ/Δ^* in chief cells does not alter stomach architecture under homeostatic conditions. Left: *Yap^+/+^; Taz^+/+^* mouse stomach; Middle: one week chasing after *Yap^Δ/Δ^; Taz^Δ/Δ^*; right: eight weeks chasing after *Yap^Δ/Δ^; Taz^Δ/Δ^*. Scale bar, 100 µm. (C) H&E images show that *Yap^Δ/Δ^; Taz^Δ/Δ^* in chief cells increases gastric gland base drop out in paligenosis. Left: control mouse stomach, 48 h after first HDTAM treatment; Right: *Yap^Δ/Δ^; Taz^Δ/Δ^* mouse stomach, 48 h after first HDTAM treatment. Scale bar, 50 µm. (D) YAP1 IHC shows the gastric gland base drop out in control and *Yap^Δ/Δ^; Taz^Δ/Δ^* mice. Top: YAP1 in control stomach, 48 h after first HDTAM treatment. Bottom: YAP1 IHC staining of YAP/TAZ deletion mouse stomach, 48 h after first HDTAM treatment. Red brackets point out the location of the chief cell zone, yellow dashed lines point out the outline of gastric glands, and red arrows point out the drop out of the gland bases. Scale bar, 50 µm. (E) Ki67 IHC highlights the *Yap^Δ/Δ^; Taz^Δ/Δ^* mice cell cycle re-entry decrease in paligenosis stage 3. Top: YAP1 in control stomach, 48 h after first HDTAM treatment. Bottom: YAP1 in YAP/TAZ deletion mouse stomach, 48 h after first HDTAM treatment. Red brackets point out the location of the chief cell zone. Scale bar, 50 µm. (F) H&E shows that *Yap^Δ/Δ^* (*Mist1^CreERT^*^2^*^/+^;ROSA26^LSL-Ai^*^9^*^/+^; Yap1^flox/flox^*, left image) or *Taz*^Δ/Δ^(*Mist1^CreERT^*^2^*^/+^; ROSA26^LSL-Ai^*^9^*^/+^; Taz^flox/flox^*, right image) in chief cells leads to increased dropout of the gastric gland base. Scale bar, 50 µm. (G) Quantification of Extended Data Fig. 2F. The control group represents mice depicted in the representative 48 h control image in Fig. 2B. (H) Ki67 IHC shows delayed cell cycle re-entry during stage 3 of paligenosis in *Yap^Δ/Δ^* mouse (middle) and *Taz^Δ/Δ^* mouse (right). Scale bar, 50 µm. (I) Quantification of mice like those depicted in panel H. The control group is from mice depicted as 72 h control population in Fig. 2D. (J) H&E (left) and Ki67 IHC (right) show that YAP1 activation in chief cells (*Mist1^CreERT^*^2^*^/+^; ROSA26^LSL-Ai^*^9^*^/+^; Nf2^flox/flox^*) is sufficient to disrupt normal gastric gland architecture and promote cell cycle entry. Scale bar, 100 µm. Data information: \*\**P* < 0.01; \*\*\**P* < 0.001 (unpaired t-test). Data are presented as mean ± SEM, derived from 10 low-power fields per condition across three independent experiments.

**Extended Data Figure 3.**
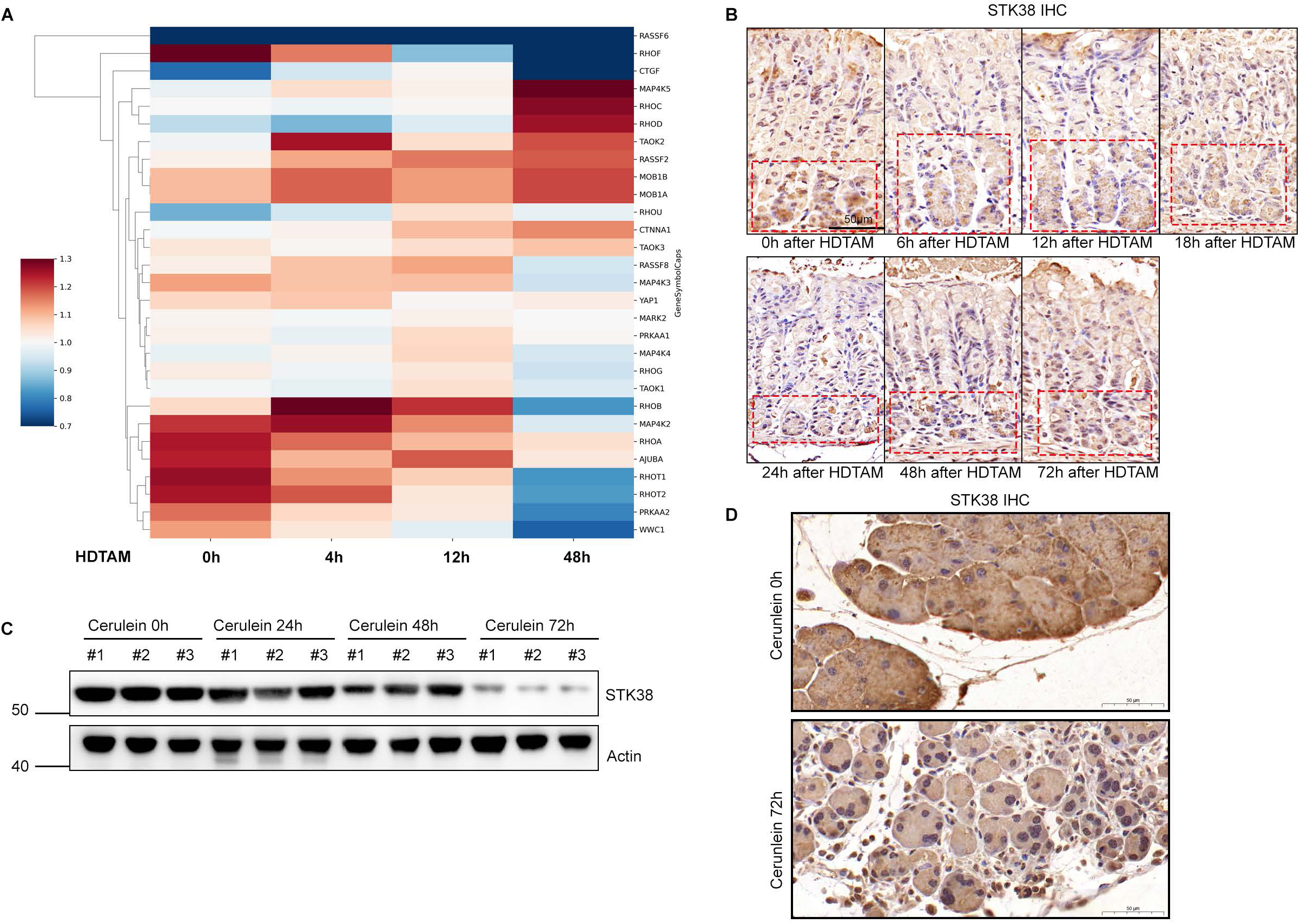
STK38 is the primary kinase regulating YAP1 activation during paligenosis. (A) Heat map showing differential expressions of a cohort of known Hippo pathway-related proteins in the stomach during paligenosis, as detected by mass spectrometry analysis. Proteins already shown in other heat maps (e.g., YAP1, STK38) are excluded (B) IHC for STK38 in C57BL/6J stomach tissue shows expression in chief cells under homeostatic conditions with decrease during paligenosis. (C) STK38 total protein abundance decreases during paligenosis in pancreas of C57BL/6J mice. (D) IHC staining of STK38 in C57BL/6J pancreas under homeostasis (top) and paligenosis (bottom).

**Extended Data Figure 4.**
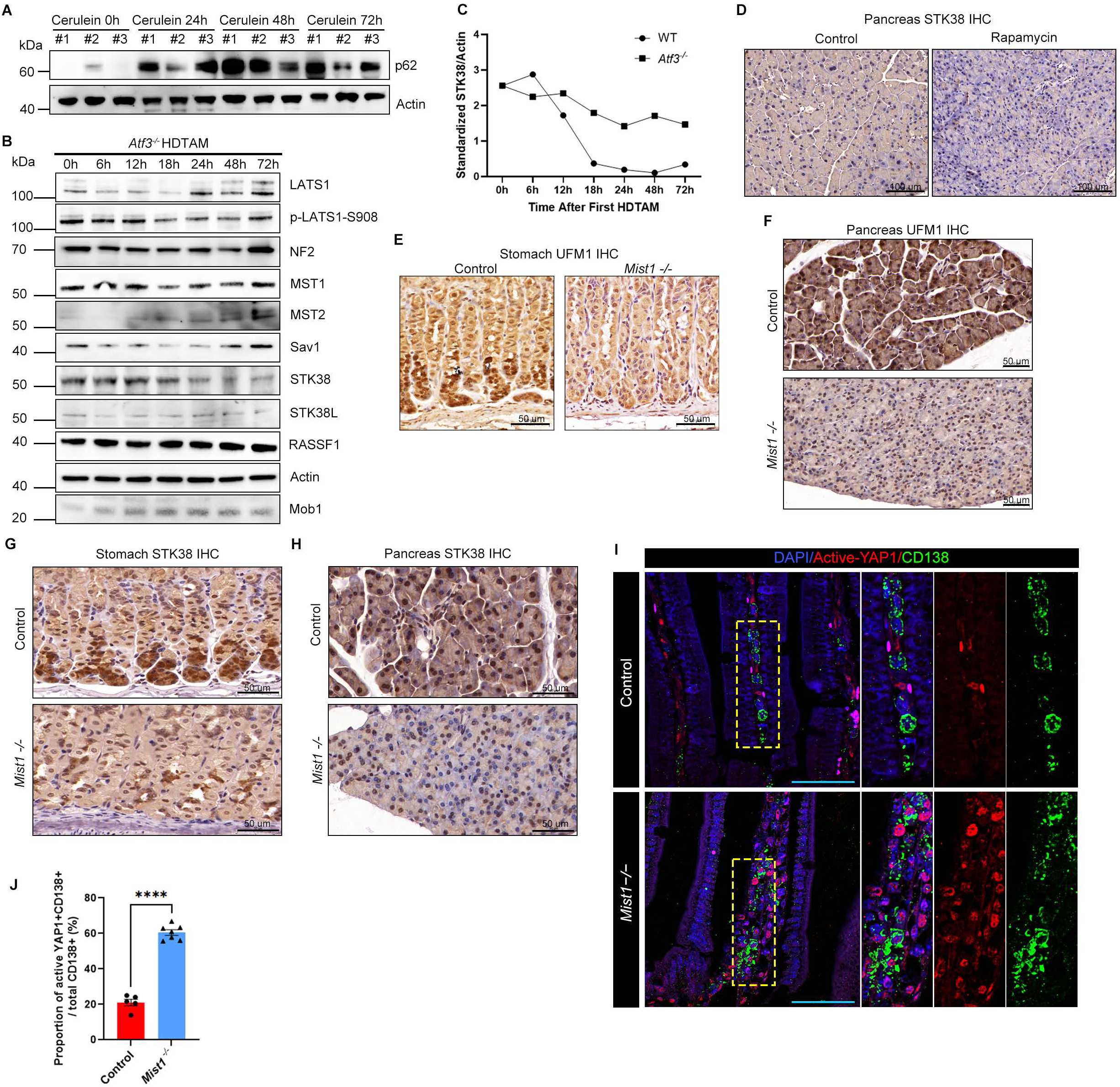
The MIST1-UFM1-STK38 axis. (A) Increased levels of active p62 are observed in the pancreas of Cerulein-treated C57BL/6J mice. (B) Total abundance of canonical Hippo pathway kinases remain unchanged in *Atf3^−/−^* mice during paligenosis, and the decrease seen in STK38 early in paligenosis is less marked with STK38 also persisting throughout paligenosis. (C) Quantification of panel B highlights how STK38 abundance declined more gradually and less markedly in *Atf3^−/−^* mice compared to wild-type mice. (D) IHC staining revealed a decrease in STK38 levels in the pancreas of rapamycin-treated mice. Scale bar, 100 µm. (E,F) IHC staining demonstrated a reduction of UMF1 in *Mist1* knockout mouse gastric glands (E) and acinar cells (F). Scale bar, 50 µm. (G,H) IHC staining illustrated a decrease in STK38 levels in the stomach (G) and pancreas (H) of *Mist1* knockout mice. Scale bar, 50 µm. (I,J) Immunofluorescence staining shows upregulation of active YAP1 in plasma cells of *Mist1* knockout mice compared to controls. The immunofluorescence staining of plasma cells (I) with subsequent statistical analysis confirms finding by (J). Scale bar, 50 µm. \*\*\*\**P* < 0.0001 (unpaired t-test). Data are presented as mean ± SEM, derived from 10 low-power fields per condition across three independent experiments.

